# Structural Basis of a Novel Heme Binding Bacterial One-Component Switch

**DOI:** 10.64898/2026.03.15.711900

**Authors:** James J. Siclari, Malvin Forson, Cullen Roeder, Eta A. Isiorho, Denize C. Favaro, Rinat R. Abzalimov, Stephen S. Gisselbrecht, Alec H. Follmer, Martha L. Bulyk, Kevin H. Gardner

**Affiliations:** Structural Biology Initiative, CUNY Advanced Science Research Center, New York, NY 10031; Ph.D. Program in Biology, The Graduate Center – City University of New York, New York, NY 10016; Ph.D. Program in Biochemistry, The Graduate Center – City University of New York, New York, NY 10016; Division of Genetics, Department of Medicine, Brigham and Women’s Hospital and Harvard Medical School, Boston, MA 02115; Department of Chemistry, University of California Davis, Davis, CA 95616, USA; Department of Pathology, Brigham and Women’s Hospital and Harvard Medical School, Boston, MA 02115; Ph.D. Programs in Biochemistry, Biology, and Chemistry, The Graduate Center – City University of New York, New York, NY 10016; Department of Chemistry and Biochemistry, City College of New York, New York, NY 10031

**Keywords:** Per-ARNT-Sim domains, Bacterial Signaling, Heme Binding Proteins, One-Component Systems

## Abstract

One-component systems (OCSs) integrate sensory and effector functions within a single protein, enabling rapid gene expression changes in response to environmental cues. Here, we characterized a novel heme binding OCS protein, FG214, from *Fimbriimonas ginsengisoli*, a redox-regulated helix-turn-helix transcription factor in which heme iron ligand state controls a monomer-to-dimer switch. Data supporting this included our observation of the FG214 PAS domain binding a hexacoordinate heme b in oxidized conditions and undergoing a slate of redox and ligand-dependent conformational changes, transitioning from a monomer to a homodimer. Spectroscopic and structural data revealed that oxidation stabilizes the likely HTH-PAS intramolecular domain interface, while reduction of the heme iron dissociates the HTH, freeing previously-sequestered homodimerization surfaces. Similar effects were seen by addition of a small molecule ferric heme ligand, as directly visualized with a 1.47 Å crystal structure of an imidazole-bound truncated construct. Using *in vitro* DNA-binding assays, we identified an artificial promoter sequence and demonstrated ligand-enhanced protein-DNA binding. Finally, we performed *in vivo* proof of concept experiments establishing FG214 as a redox-sensitive scaffold for biosensor engineering. Together, these findings define FG214 as a novel heme-binding PAS DNA binding protein, complementing known heme-PAS two-component signaling switches.

## Introduction

Bacteria rely on an array of sensory and signal transduction proteins to detect and respond to environmental fluctuations. Many such responses are mediated by one-component systems (OCS), in which the sensory and effector domains reside within a single polypeptide(1). Among these proteins, Per-ARNT-SIM (PAS) domains are often found playing sensory roles given their remarkable adaptability, binding diverse small molecules to allosterically regulate partner domains differentially(1–4).

A subset of bacterial PAS domains coordinate heme prosthetic groups, enabling direct sensing of gases, redox potential, and metabolic state(5). Much of our understanding of this regulation stems from one such type of system, the *Bradyrhizobia* FixL histidine kinase-FixJ response regulator pair, where changes in O_2_ occupancy at the distal position alters net kinase activity of FixL, along with subsequent FixJ phosphorylation and transcriptional activation. However, questions remain open about both FixL signaling – particularly how signals are transduced from the hemes to activate this enzyme – and more broadly about their applicability to other heme-PAS proteins(6–9).

Beyond their physiological roles, PAS-containing OCS proteins offer powerful platforms for biosensing and synthetic regulation due to their compact architecture and modular signaling logic. The blue-light responsive activator EL222, for example, undergoes a conformational switch upon illumination that releases its helix-turn-helix (HTH) effector, activating transcription(10–13). Previous work from our lab and others have engineered this system for uses in prokaryotic and eukaryotic model systems(14–20). Similar sensory logic could, in principle, be harnessed for small molecule or redox driven systems.

Here we describe FG214, a HTH-PAS transcription factor from *Fimbriimonas ginsengisoli*(21), that coordinates a heme b within its PAS domain via bis-histidine ligation of the ferric iron as purified *in vitro*. We show that redox changes or exogenous imidazole induce structural rearrangements which convert FG214 from a monomer into a homodimer, enabling DNA binding. Our work identifies a novel one-component heme binding PAS protein and establishes a mechanistic framework for redox-regulated “effector release” while also providing proof of concept experiments for its utility as an *in vivo* biosensor. These findings expand the functional landscape of PAS domains and suggest new strategies for engineering redox-sensitive systems.

## Results

### Redox-dependent global conformational changes

FG214 was identified in a bioinformatics screen for PAS domain-containing transcription factors with potential chemosensory triggers(22). These analyses predicted an N-terminal LuxR-type DNA-binding tetrahelical helix-turn-helix (HTH) DNA-binding domain and a C-terminal PAS domain, connected by a predicted helical linker. While these two types of domains are commonly found together (over 6000 occurrences in SMART(23) as of July 2026), the vast majority of these have a domain architecture with an N-terminal PAS sensor and C-terminal LuxR HTH DNA-binding domain, as previously seen in EL222(10) (**Fig. 1a,b**). Following overexpression and purification from *E. coli*, FG214 was visibly red, consistent with metal binding. Subsequent UV-visible absorbance spectroscopy and LC-MS revealed a heme b ligand(24) (**Fig. 1c,d,e**), and redox titrations demonstrated reversible spectral changes consistent with a redox-active heme(25–27) (**Fig. 1d**).

**Figure 1:**
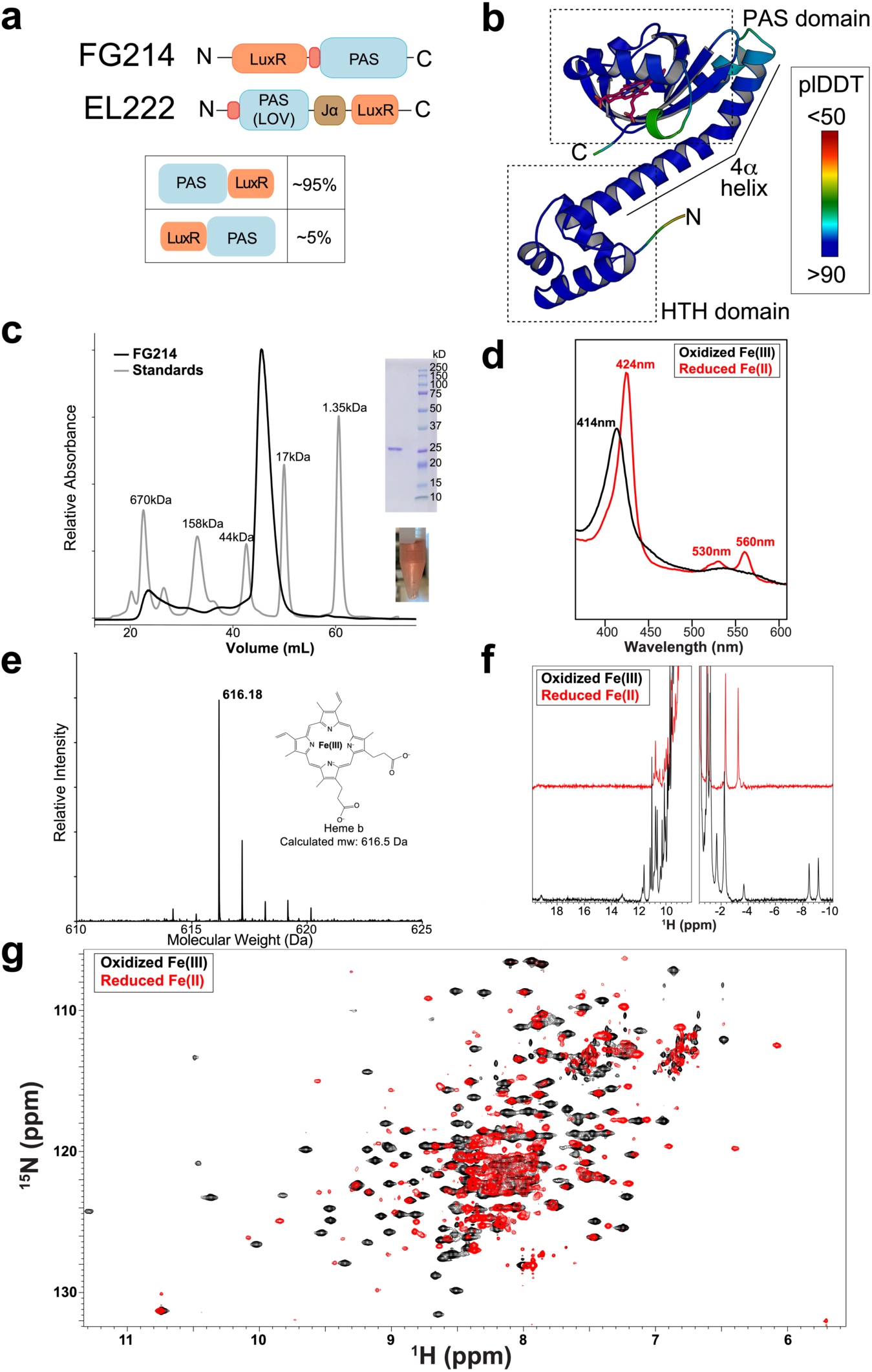
FG214 is a redox-sensitive heme-binding protein. **a)** Domain architecture of FG214 and another PAS-containing transcription factor, EL222. **b)** AlphaFold3 model of FG214, highlighting locations of HTH and PAS domains, as well as the connecting 4α helix. **c)** Superdex S200 SEC chromatogram of FG214 compared to standards of known molecular weight. (inset) SDS-PAGE gel verifying expected MW (24 kDa predicted from sequence) and image of purified protein. **d)** UV-visible absorbance spectra of oxidized (black) and reduced (red) FG214. **e)** Electrospray ionization LC-MS analysis confirming the presence of heme b within FG214 isolated from *E. coli*. **f)** ^1^H NMR spectra of oxidized (black) and reduced (red) FG214. **g)** ^15^N/^1^H TROSY spectra of oxidized (black) and reduced (red) FG214.

Reduction of the heme iron induced widespread structural changes detected by ^1^H and ^15^N/^1^H TROSY NMR (**Fig. 1f,g**). In the oxidized state, the spectra exhibited well-dispersed resonances of uniform intensity, suggesting a folded protein. We observed ^1^H peaks of oxidized FG214 distinctively upfield of 0 ppm, likely arising from protons interacting with a low-spin ferric Fe(III) state in the heme(28). These peaks were greatly perturbed by reduction, typical of a paramagnetic Fe(III) to diamagnetic Fe(II) spin system transition(29).

More globally, reduction caused extensive chemical shift perturbations and peak broadening in FG214 ^15^N/^1^H TROSY spectra, suggestive of a large-scale protein conformational change. While we lack the chemical shift assignments needed to fully establish the nature of this event using solely solution NMR, we can establish two key features by comparing spectra acquired from a series of FG214 truncations (1-214, 73-214 [Δ72], 87-214 [Δ86]) (**Fig. S1**). Several peaks in this series that are present in all three spectra – and hence must arise from residues in the PAS domain or an N-terminal 25 residue section – show a progressive linear change in chemical shift, strongly suggesting that the truncations alter an equilibrium. Coupled with AlphaFold3 models of the variants, this pattern was highly reminiscent of the light-triggered helix release equilibrium seen in the photosensitive *As*LOV2 system with an FMN-bound PAS domain(30, 31). These same peaks exhibited reduction-triggered chemical shifts as well, strongly suggesting that a heme-based mechanism also alters this equilibrium *in vitro* (**Fig. S1**). Taken together, our solution NMR data indicate substantial redox-triggered changes in FG214 structure suggestive of a triggered release of N-terminal regions before the PAS domain.

### Reduction promotes release of the HTH domain

Hydrogen-deuterium exchange mass spectrometry (HDX-MS) was used to elucidate how heme reduction shifts the conformational landscape of FG214. HDX-MS protection patterns of both the oxidized and reduced form of the protein are generally consistent with AlphaFold3 predictions, including placements of all 2° structures in both domains. The predicted long 4α helix is also visible in these protection patterns, but notably, it shows a marked increase in deuteration levels upon reduction (**Fig. 2a,b,c**). This interdomain 4α helix appears to make direct contact with the FG214 PAS domain in the oxidized monomer AlphaFold3 model. Our experimental data clearly suggests reduction destabilizes the 4α helix from the PAS core, demonstrating allosteric release of the effector domain. This is a mechanism seen multiple times previously in prokaryotic and eukaryotic signaling proteins(31–34). We also see reduction influenced changes in the PAS domain around the known ligand binding regions, mainly in the Eα helix and Hβ strand, consistent with a redox-triggered shift in the geometry or volume of the heme binding pocket. These features suggest that the oxidized state adopts a stable monomeric “off” conformation, whereas reduction destabilizes the fold.

**Figure 2:**
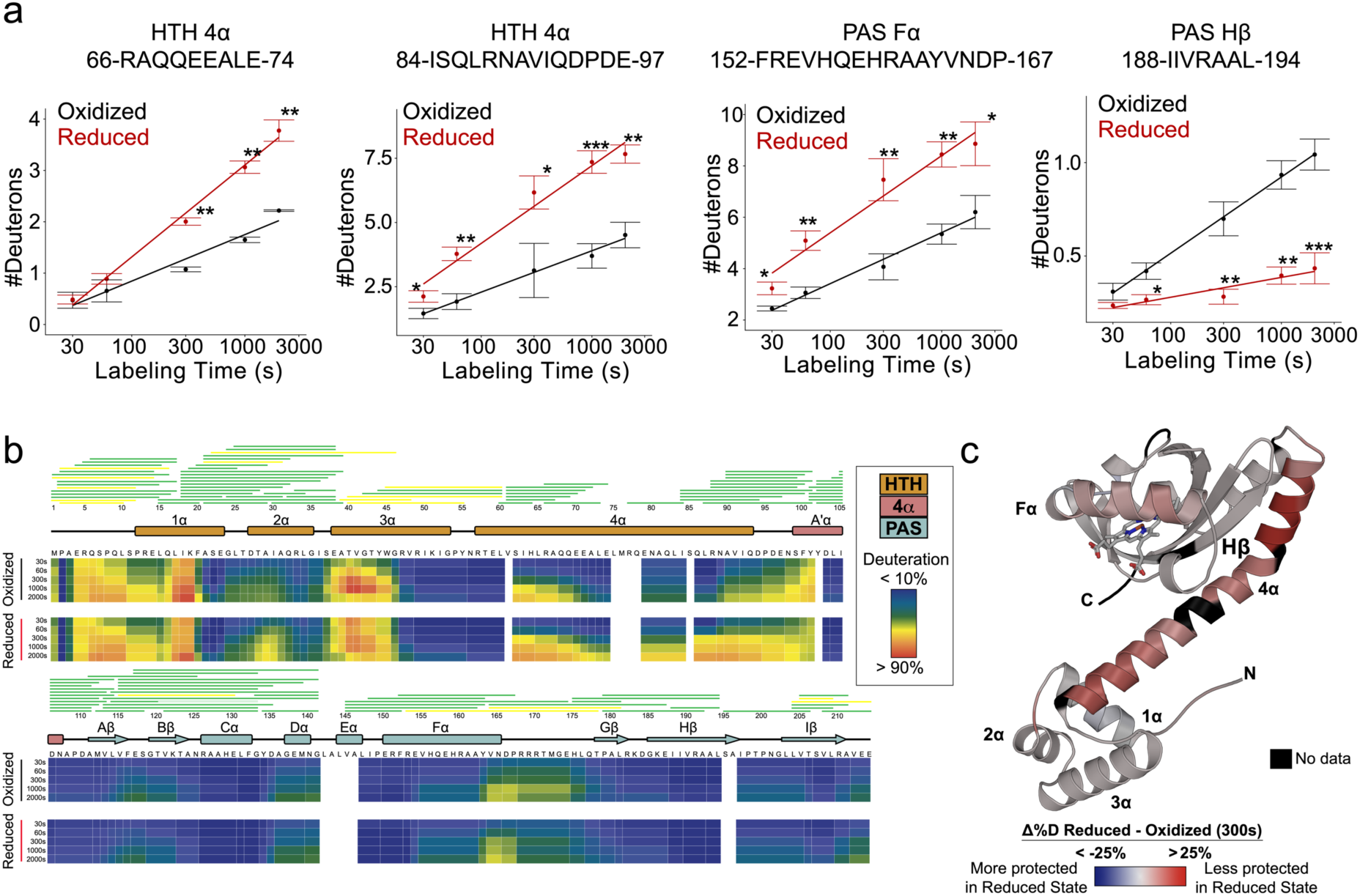
HDX-MS indicates that FG214 employs an effector-release mediated mode of activation. **a)** Peptide deuteration uptake plots of selected regions of FG214 under oxidized (black) or reduced (red) conditions. **b)** Heat map showing percent change in deuteration over time between oxidized and reduced states. Secondary structure diagram shows HTH domain in orange, Aʹα in pink, and PAS domain in cyan. **c)** Heat map of reduction-triggered changes in HDX protection as data mapped onto FG214 AlphaFold3 model. Red color indicates less protection in reduced form; blue indicates more protection in reduced form; black indicates no data due to gaps in peptide coverage. Asterisks denote statistical significance derived from a two-tailed Welch’s t-test.

### Heme coordination chemistry is linked to FG214 dimerization

The iron in heme b is typically coordinated by a combination of histidine and/or methionine residues in the oxidized state(35). Identifying the coordinating residues within FG214 allows for targeted mutagenesis to probe structure-function relationships and potentially shift the equilibrium between inactive and active conformations. Guided by AlphaFold3 structural predictions (**Fig. 3a**), we identified four candidate residues (His 156, His 159, Met 172, His 175) and sought to experimentally determine their roles in heme coordination.

**Figure 3:**
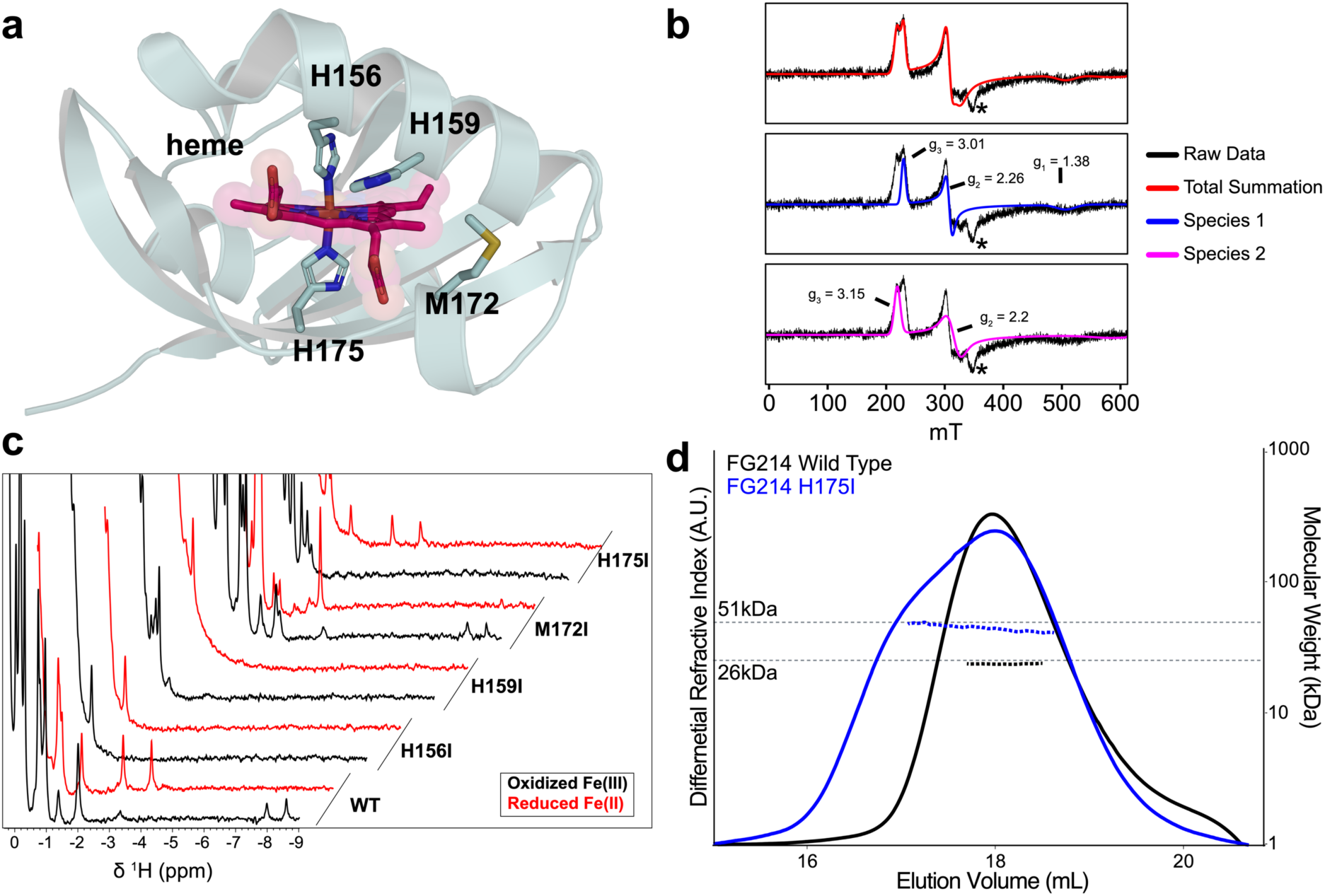
FG214 binds heme with two histidine residues in the oxidized state. **a)** AlphaFold3 model of full length FG214 heme binding pocket. **b)** X-band EPR spectra of oxidized FG214 showing two different low spin ferric states. Asterisk indicates a small background Cu signal. **c)** ^1^H NMR spectra of oxidized (black) and reduced (red) wild type (bottom, reproduced from Figure 1f) and point mutants of potential liganding residues. **d)** SEC-MALS using a Superdex 200 column of WT (black) and H175I (blue) FG214 proteins.

As an initial examination, we acquired X-band electron paramagnetic resonance (EPR) spectra of oxidized FG214, revealing two rhombic S=1/2 populations (**Fig 3b**). Both species had g-values supporting the assignment of 6-coordinate low spin-ferric heme, sometimes referred to as highly anisotropic/axial low-spin (HALS) systems(36). The first, less rhombic species exhibited g-values (3.01, 2.26, 1.38) similar to many other HALS proteins where the heme cofactors are bound by two histidine residues arranged in an antiparallel geometry (**Fig. 3b**), consistent with our ^1^H NMR spectra displaying upfield-shifted resonances (**Fig 1f**). The second species is more rhombic and only two g-values are observed in this magnetic field range (g-values: 3.15, 2.2). While we cannot rule out that this second species arises from a second orientation of the bis-His coordination, it is also possible that this more rhombic population indicates a potential His-Met coordination in the oxidized state with similar g-values observed for these systems(36).

To further test this model, we generated a series of single point mutations targeting the four histidine and methionine residues predicted in the binding pocket and substituting each with isoleucine to maintain hydrophobicity and approximate sidechain volume. We used UV-visible absorbance spectroscopy to compare relative heme loading across mutants, using samples with matched protein concentrations (as judged by A_280 v_alues) and, focusing on changes in the Q-and Soret band regions. While M172I exhibited near-wildtype ability to bind heme as judged by the Q-and Soret band peak intensities, the other three mutants were impaired. This was most evident with the H156I mutant, particularly with the near absence of a Soret band in the reduced state (**Fig. S2, Table S1**). The H159I mutant also exhibited a distinct spectral signature, including a new absorbance maximum at 662 nm and an altered color in solution, reflecting perturbed coordination geometry (**Fig. S2, Table S1).** The persistence of partial binding among all mutants suggests compensatory heme coordination, underscoring some degree of pliability in the FG214 heme binding site.

^1^H NMR spectra and size exclusion chromatography of the mutated proteins further revealed structural changes caused by these substitutions. Focusing first on the NMR signals upfield of 0 ppm, H156I and H159I displayed substantial peak losses in this region, consistent with global destabilization of the protein (**Fig. 3c**). M172I and H175I were subtler in their changes, but notably with oxidation-specific effects, with M172I being most strongly perturbed in the reduced form and H175I in the oxidized state. Coupled with UV-visible absorbance data and the AlphaFold3 prediction, we assign His 156 (proximal) and His 175 (distal) as the heme-coordinating residues, with some potential role for Met 172 as an alternative distal participant.

To explore the roles of these heme-coordinating residues in the FG214 quaternary structure, we used size-exclusion chromatography with inline light scattering (SEC-MALS) to characterize changes in solution shape and mass. While the lack of heme binding by H156I precluded us from acquiring meaningful SEC-MALS data on this variant, we obtained data clearly showing that wildtype FG214 is monomeric in solution and H175I is markedly shifted towards a dimeric species (**Fig. 3d**). Taken together, our solution data clearly support a model where changes near the heme site affect both an intramolecular HTH-PAS interaction predicted in the oxidized state, leading to protein dimerization.

### Crystal structure of the activated dimer

Guided by truncation analysis (**Fig. S1**), we successfully crystallized an imidazole-bound construct lacking the first N terminal 72 residues (Δ72), encompassing part of the HTH 4α helix through the PAS domain. In solution, this construct shares many features in common with the full-length protein, including being monomeric by SEC-MALS when oxidized, having nearly identical UV-visible absorbance characteristics, and similar ^1^H NMR shifts – with a notable shift in the location of heme vinyl chemical shifts near –8 ppm, showing that changes outside the PAS domain impact the environment near the heme (**Fig. S3**). During crystallization trials for FG214, we observed that addition of imidazole was required to obtain crystals, which we attribute to imidazole’s well-characterized ability to ligand ferric heme proteins. In doing so, it is known to bind the distal coordination site of heme, outcompeting native protein sidechain interactions, reproducing the electrostatic properties of Fe(II)-O_2 c_omplexes(37, 38).

From these imidazole-bound FG214(Δ72) crystals, we obtained a 1.47 Å structure of an imidazole-bound homodimer (**Fig. 4a**), mediated by both HTH-HTH and PAS-PAS interactions. Within the PAS domain, H156 serves as the proximal heme-coordinating residue, while an imidazole molecule occupies the distal site, fully corroborating our spectroscopic and mutagenesis data. Meanwhile, the H175 sidechain, which we expect to normally serve as the distal ligand, was displaced into an unstructured loop that meets the other monomer at the dimerization interface. Of note, the nearby M172 sidechain is similarly outside of the PAS domain in one of the two chains but is not resolved in the other. These structural changes clearly implicate linkage between heme coordination and protein oligomerization state together.

**Figure 4:**
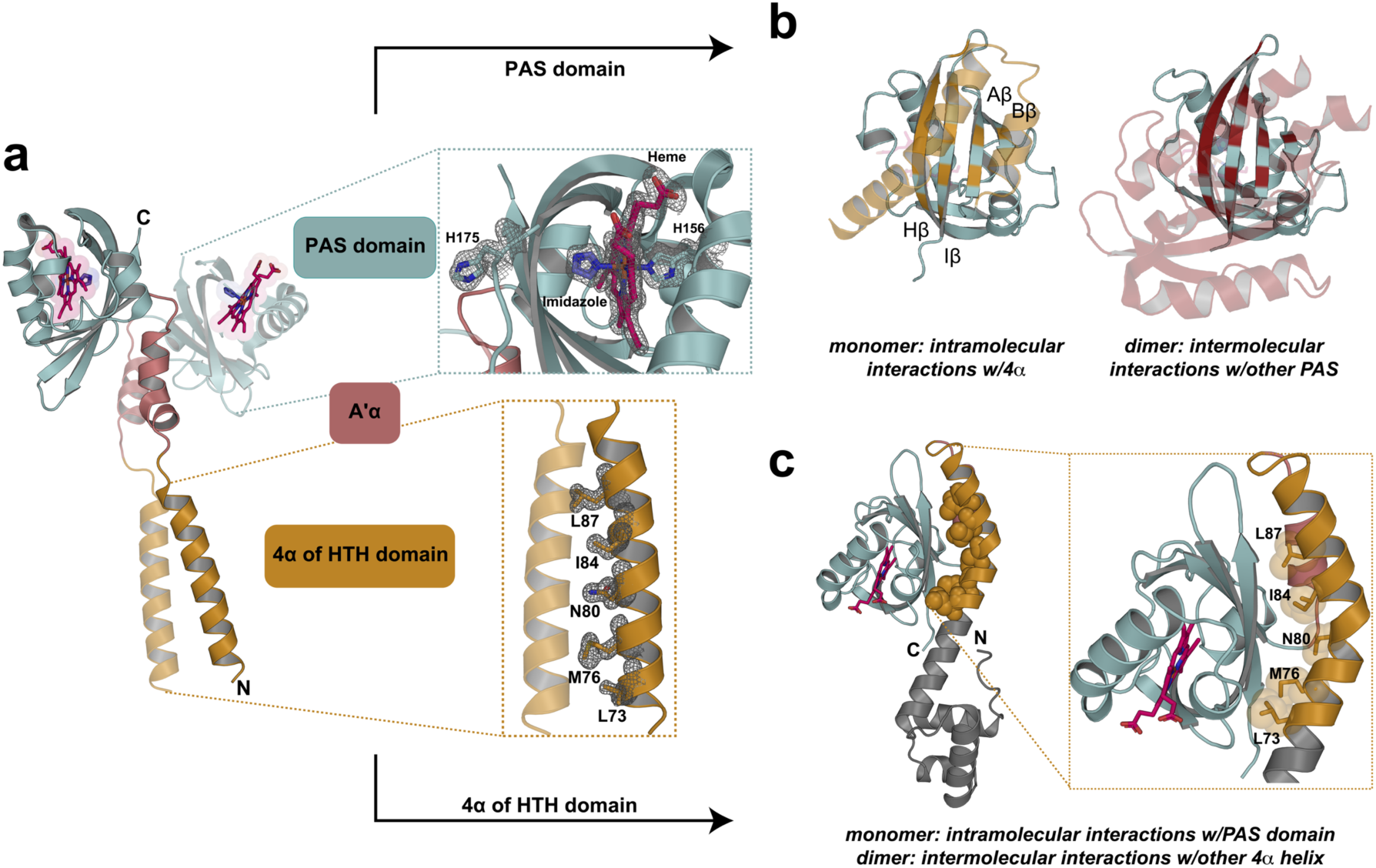
Crystal structure of an N-terminally truncated FG214 protein reveals imidazole driven dimerization. **a)** 1.47 Å crystal structure of FG214(Δ72), PDB:10JX. Mesh shows electron density. Insets show heme binding cavity and hydrophobic interface of HTH 4α helix. **b)** FG214 PAS domain highlighting residues 4 Å away from the 4α helix in the AF3 monomer model (left) and 4 Å from the other PAS domain in the dimeric crystal structure (right). **c)** AlphaFold3 model of oxidized full length FG214 highlighting the locations of the 4α residues involved in the dimeric structure of Figure 4a.

Examining the FG214 dimer, we saw potential interactions between the two chains mediated by both PAS/PAS and 4α/4α contacts. At the PAS domains, these were chiefly involving Aʹα helix/β-sheet and β-sheet/β-sheet interactions; of note, the β-sheet residues are also predicted by AlphaFold3 to make an intramolecular monomeric interface with 4α (**Fig. 4b**). At the HTH domains, hydrophobic sidechains of the two 4α helices interact to form the dimer; reminiscent of coiled-coil interfaces of transcription factors(39–41). And again, the 4α residues at the dimer interface are predicted to play an integral role in stabilizing the monomeric form (**Fig. 4c**).

Soaking these same crystals with sodium dithionite (DT) yielded datasets leading to a 1.67 Å structure of FG214(Δ72), revealing a homodimer with nearly identical 4° structural arrangements as previously seen. However, there are distinct changes at the heme – which has a His 156-Met 172 iron coordination pair and His 175 on an extended loop pointing into solution (**Fig. S4**). This raises the potential that alternative His-His and His-Met heme coordination pairs – as suggested by our EPR and mutagenesis results in the full length protein (**Fig. 3**) – may exist and be biased in different crystal forms of FG214 variants. Indeed, AlphaFold3 modeling suggests that removal of the HTH domain favors His-Met coordination in the monomer (**Fig. S5**), perhaps due to altered states of the outer β-sheets within the PAS core due to the shortened 4α helix. Regardless, these structural data strongly implicate changes at the heme site being amplified by the surrounding protein to affect the monomer:dimer equilibrium of FG214, as is commonly seen as a regulatory mechanism in other proteins.

### Identification of an artificial DNA binding site

To explore the functional importance of these changes, we aimed to identify a suitable artificial DNA sequence for FG214 binding, both to enable tests of our hypothesis of imidazole-driven activation and inform downstream functional assays and future bioengineering. To do so, we used universal protein binding microarrays (PBMs) to screen FG214 for *in vitro* sequence-specific DNA binding activity against all possible8 bp sequences(42, 43). Analysis of PBM-derived scores from these experiments of these data yielded a DNA binding specificity motif for FG214 (**Fig. 5a**), with a notable bias towards a purine-rich strand on one side of an 8 bp sequence.

**Figure 5:**
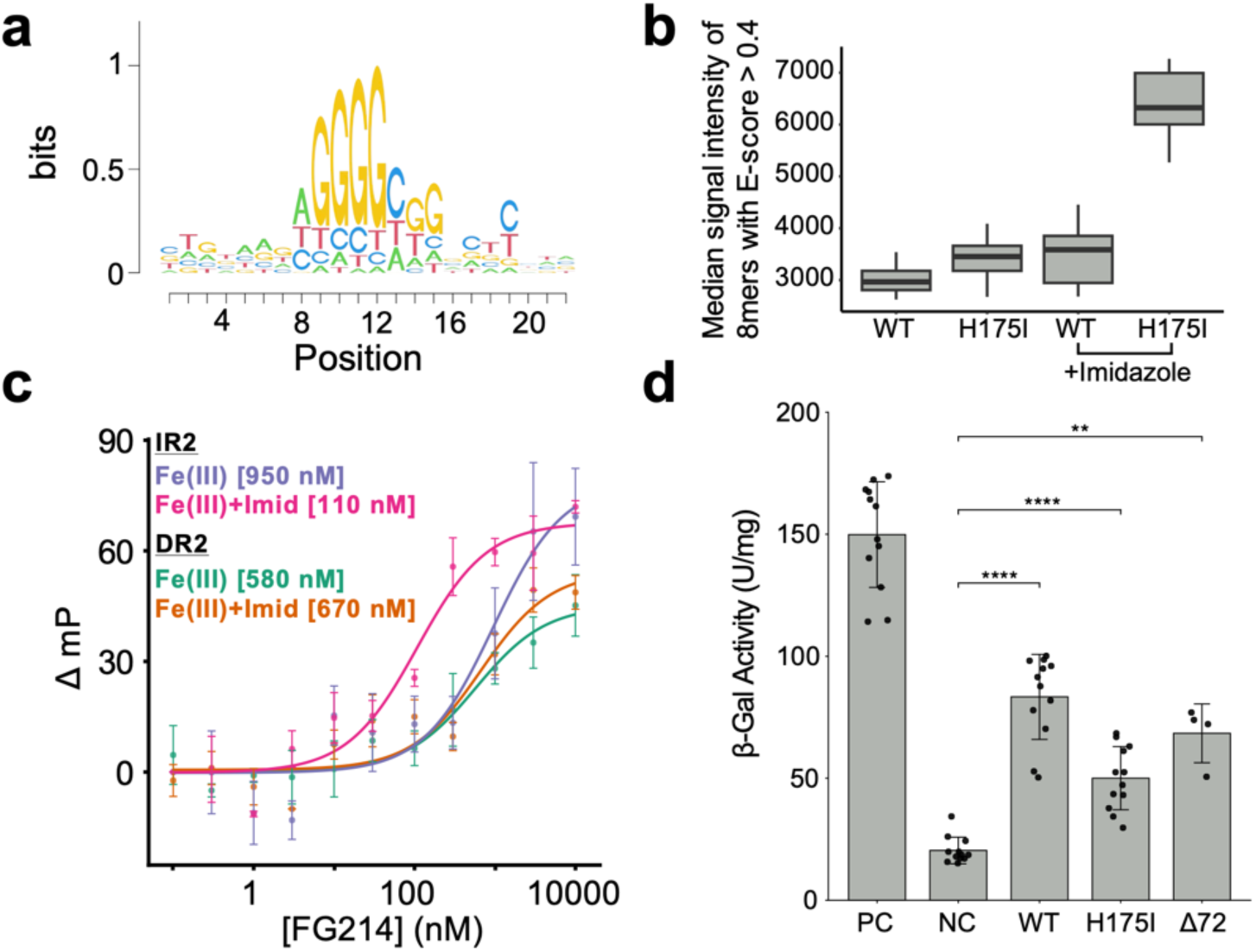
FG214 binds a GC-rich palindromic promoter region enhanced by imidazole in the ferric state. **a)** DNA half-site for FG214 binding identified by universal protein-binding microarray (PBM). **b)** Median intensity of specifically-bound 8-mers in PBM for FG214 WT and H175I in the absence and presence of 500 mM imidazole. **c)** Fluorescence polarization measurements of 1 nM FAM-labeled DR2 or IR2 sequences with increasing concentrations of FG214 in the absence and presence of 10 mM imidazole. All individual measurements were collected in triplicate; points shown are average ± 1 standard deviation**. d)** β-galactosidase activity quantified following bacterial two hybrid (BTH) analysis of split adenylyl cyclase vectors (negative control, NC) or fused to leucine zippers (positive control, PC), FG214 full-length, FG214 full-lengthH175I, or FG214 (Δ72). Statistical significance derived from a Welch’s t-test, with **** = p < 0.0001 and ** = p < 0.01. Individual datapoints are shown, with top bar drawn at mean value and error bars showing ± one standard deviation from the mean.

Given the role of the distal His 175 in maintaining the FG214 monomer, we also tested whether ligand-induced displacement of this sidechain in a wild-type background could promote DNA binding. Analysis of PBM fluorescence intensities across the array revealed enhanced binding by the H175I mutant relative to wild-type FG214, with further increases for both constructs in the presence of 500 mM imidazole (**Fig. 5b**). These observations suggest that imidazole-induced displacement of His 175 facilitates domain rearrangements sufficient for dimer-mediated DNA binding, thereby partially activating FG214 under ferric conditions. These findings align with prior studies showing that exogenous imidazole can act as a functional mimic for *in vivo* heme ligands(44–46). In this context, imidazole binding appears to trigger an active conformation of FG214 without the need for heme reduction.

To develop an optimal high-affinity DNA binding site for *in vitro* assays, we constructed a series of DNA sequences containing direct or inverted repeats of the PBM-identified motif (GGGGCGGGG), assuming that this sequence is a likely half-site for binding a FG214 dimer. We generated 14 FAM labeled binding substrates with varying half-site orientation and spacer length from 1-7 bp (**Fig. S6**). Each DNA construct was labeled with FAM (fluorescein derivative) on the 5ʹ end to enable quantitative binding analysis via fluorescence polarization/anisotropy. Oligos containing direct repeats of the motif bound FG214 weakly and non-specifically, and without any dependence on mutation or imidazole concentration (**Fig. S6**). In contrast, oligos with inverted repeats of the motif displayed stronger binding, particularly for the H175I variant, and often with imidazole dependence (**Fig. S6**). From these experiments, a particularly clear exemplar of FG214 DNA binding characteristics are evident from oligos with a 2 bp central spacer: Inverted copies of this motif (IR2; **Table S3**) exhibited the most pronounced imidazole-dependent binding response (without imidazole: K_d_ ≈ 950 nM; with 10 mM imidazole, affinity increased by approximately an order of magnitude to an apparent K_d_ ≈ 100 nM) (**Fig. 5c**). In contrast, direct repeats around the same 2 bp spacer (DR2; **Table S3**) showed moderate binding of K_d_ ≈ 600 nM with no imidazole dependence.

### *In Vivo* Validation of FG214 homodimerization

To explore the potential for FG214 to adopt the homodimeric state that appears to be required for DNA binding, we employed a bacterial two hybrid (BTH) assay to probe FG214 homodimerization in *E. coli.* By fusing FG214 (full length, mutant, or the Δ72 truncation) to fragments of a split adenylyl cyclase we can analyze FG214 homodimerization following co-transforming into *E. coli* lacking its native adenylyl cyclase(47, 48). Here, complementation of the split cyclase will only occur with sufficient FG214 homodimerization, facilitating cyclic AMP (cAMP) production from ATP and activating CAP-dependent expression of a β-galactosidase which can be easily assayed on plates or in solution as a proxy for dimerization. Employing this BTH assay by quantitatively measuring β-galactosidase activity in cell lysates in the presence of ortho-nitrophenyl-β-D-galactopyranoside (ONPG), we saw significant increases in β-galactosidase activity in all three FG214 constructs, confirming of FG214 homodimerization in *E. coli* grown under standard aerobic conditions (**Fig. 5d**). Taken together, these cell-based data are all consistent with our *in vitro* biochemical and structural results.

## Discussion

Our work provides structural and mechanistic insight into a heme-binding PAS-HTH one-component DNA-binding protein, FG214. By characterizing the monomeric inactive oxidized state, the intermediate reduced state, and the active DNA-bound homodimer triggered by imidazole, we define three likely signaling states of the protein (**Fig. 6**). Together, these data support a model in which FG214 employs an effector-release activation mechanism to sense and respond to environmental gases, a framework previously observed in other PAS-containing signaling systems. In addition to expanding our knowledge of bacterial signaling proteins, we have laid the foundation for the development of FG214-based transcriptional reporter systems.

**Figure 6:**
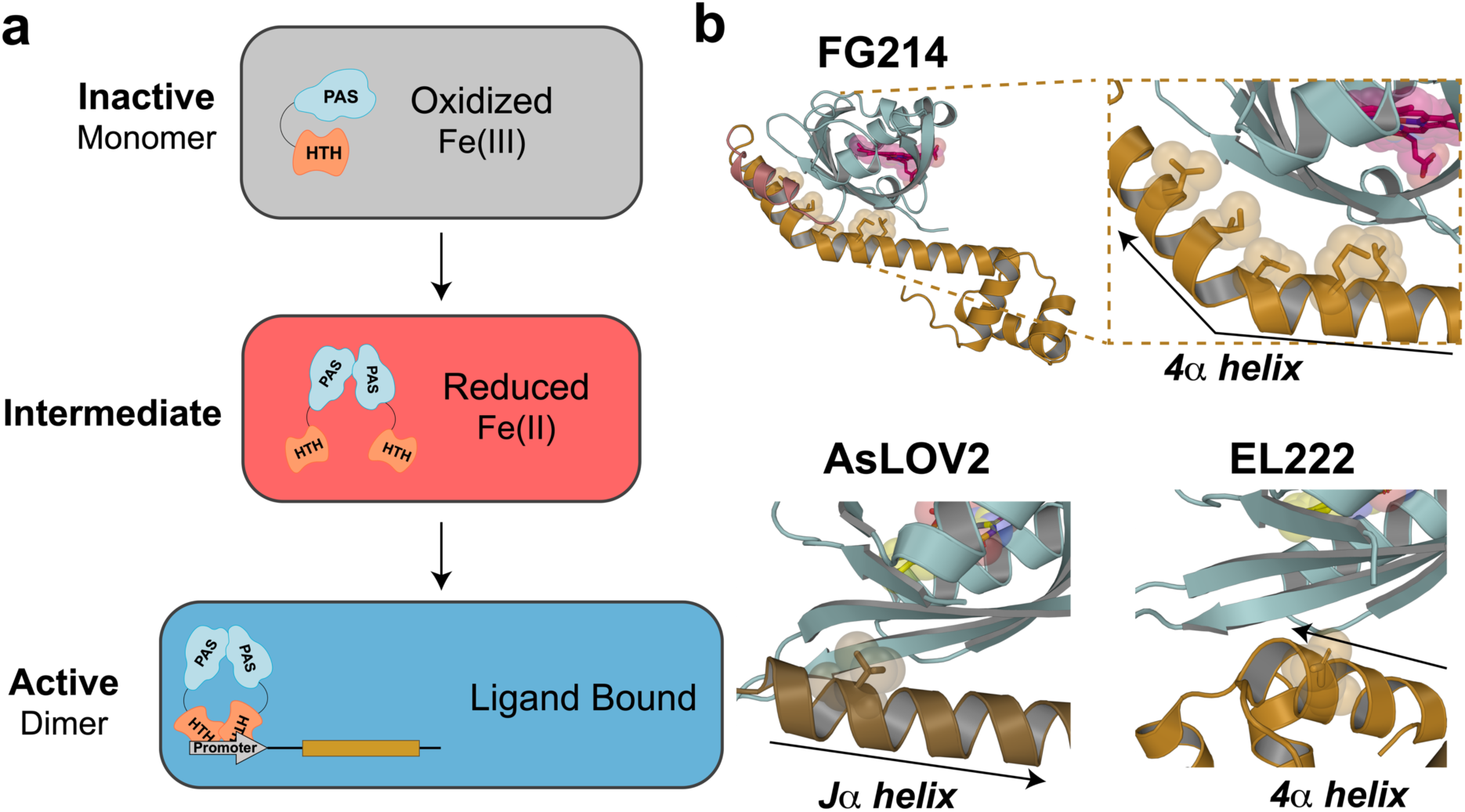
Proposed signaling model for FG214, with a signal-induced monomer to dimer transition activating DNA binding, like other PAS domain sensors. **a)** Proposed schematic of FG214 activation. **b)** EL222 and AsLOV2, PAS domain containing proteins that rely on interactions between auxiliary helices and β-sheet surfaces to maintain inactive states. Note that auxiliary helices can originate either N-or C-terminal of the PAS domain and run in either direction across PAS β-sheet surface.

### Effector-release mechanism

Structural and dynamic analyses of the oxidized FG214 monomer validate the AlphaFold prediction for the HTH 4α helix to form key contacts with the PAS domain β-sheet, particularly via HDX-MS analysis showing this helix to be well ordered. This conformation masks HTH and PAS surfaces required for dimerization, as demonstrated by our solution measurements. Upon Fe(III) reduction or ligand binding, however, this helix is markedly destabilized, demonstrating that 4α helix displacement is required for dimerization, as shown by the constitutively dimeric H175I variant and the crystal structures of the activated homodimer. Indeed, our crystal structures of reduced and imidazole-bound FG214(Δ78) confirm that the newly-freed PAS β-sheet and HTH 4α helix contribute substantial homodimer interfaces, supporting a model in which allosteric changes initiated at the heme cofactor reorganize nearby FG214 sheet/helix interactions as a key trigger to transition from a monomeric “off” state to a dimeric DNA-binding “on” state.

This activation mechanism parallels those used by certain other PAS sensors, including the light-activated AsLOV2 and EL222 from the LOV subfamily(10, 12, 30) (**Fig. 6**). In those cases, allosteric signals triggered by illumination or redox changes at an internal flavin cofactor relay across the LOV β-sheet, displacing the Jα helix (AsLOV2(30, 31)) or the 4α helix (EL222(10)) as part of the activation process. We underscore that the helices involved in these stimuli-dependent binding/unbinding equilibria bind to equivalent spots on the PAS β-sheet, despite coming from very different locations in the primary sequence compared to the PAS domain itself – from immediately C-terminal (AsLOV2 Jα) to 70-80 residues C-terminal (EL222 4α) to 20-40 residues *N-terminal* (FG214 4α) – and binding in different orientations (**Fig. 6b**). This suggests evolutionary conservation of a common signaling mechanism despite marked differences as substantial as the order of domains(49, 50). Upon freeing the 4α helix, the FG214 β-sheet becomes accessible for homodimerization, utilizing the same residues just liberated to do so (**Fig. 4b,c**). This is very reminiscent of the proposed signaling mechanism for EL222(10, 12, 13), despite differences in the controlling cofactor (heme b vs. FMN) and domain orientation (HTH-PAS vs. PAS-HTH).

### DNA binding specificity and implications for dimerization

Our DNA-binding data across constructs with varying spacer lengths provide key information into the monomer-dimer equilibrium of FG214 as well as functional data of the active state. We interpret the binding to direct-repeat sequences – relatively weak, relatively low change in fluorescence polarization, and chiefly unaffected by changes in spacer, addition of imidazole or the H175I mutation – likely reflects low-affinity recruitment of monomeric FG214 to individual half-sites. In contrast, inverted repeats displayed higher affinity, more stimulus dependence, and more variation by spacer length. Coupled with the higher ΔmP for the saturated inverted repeats when compared to the direct, these data are consistent with engagement by an activated dimeric form of FG214. Given that GC rich regions are implicated in transcriptional control of oxidative stress genes(51–53) and homologs of IR2 can be found throughout the *F. ginsengisoli* genome upstream of many redox dependent ATP-binding cassette (ABC) transporters(54, 55), we speculate that FG214 may have an *in vivo* role controlling the expression of these genes, but this remains to be experimentally validated, although the propensity for FG214 to dimerize in an *in vivo* environment supports involvement in redox associated pathways. While such cellular experiments are outside the scope of this work, we believe that the correlation we see between *in vitro* and *E. coli* dimerization of FG214 truncation or the H175I mutation – a step required for DNA binding – provides both useful reagents and frameworks for those studies.

### Comparison with other heme-binding PAS sensors

Heme-binding PAS domains have been previously reported in other proteins, but never in an OCS context. Of these, several gas-sensing bacterial PAS proteins have been best studied, including FixL, Aer2, and EcDos/DosP(56–59). The gas binding abilities of hemoproteins also make them attractive bioengineering tools for treatment of carbon monoxide, as is the case with RcoM, another heme binding PAS OCS(60, 61). Other proteins, like the FlrB histidine kinase, use the PAS fold to detect levels of heme directly as opposed to using heme as a prosthetic group to sense secondary ligands, as bacteria often sequester heme from symbionts or host organisms as an iron source(62–64).

Among known heme-binding PAS proteins, the oxygen-sensing kinase FixL remains the best characterized and is often regarded as the canonical model. In *Bradyrhizobium japonicum*, FixL forms part of a two-component system in which the heme-binding PAS domain modulates kinase activity in response to oxygen levels. In its oxidized state, FixL binds heme in a pentacoordinate geometry through a single proximal histidine(46, 65, 66). Researchers also used ligands like imidazole or cyanide to mimic O_2_ binding under oxidized conditions *in vitro*(56, 67). Such binding events reshape a preformed PAS homodimer within FixL(68), leading to conformational changes within a coiled-coil linker to a downstream histidine kinase, controlling enzymatic activity as it does with minor changes in higher order structure(69).

In contrast, FG214 coordinates heme via a chiefly bis-histidine ligation in the ferric state and undergoes distal coordination rearrangement upon activation. Subsequently triggered protein conformational changes are also distinctly different, as FG214 uses heme sensing to control a monomer:dimer equilibrium for activation. The contrast between pentacoordinate proximal-His sensing in FixL and hexacoordinate bis-histidine distal-coordination switching in FG214 illustrates the mechanistic diversity within the heme-PAS superfamily to couple similar environmental cues and relay them into downstream signaling. The comparison underscores the remarkable modularity and adaptability of PAS domains across species and regulatory contexts.

### Implications for redox signaling and synthetic applications

By uncovering the molecular basis of heme-based sensing in FG214, we have revealed how structural modularity of PAS domains within one-component systems enables evolutionary diversification of signaling logic to respond to different signals with different domain orientations while retaining some mechanistic similarity with effector release. FG214 thus represents a new prototype for heme-regulated transcriptional switches, expanding the known repertoire of OCS PAS signaling mechanisms beyond light and ligand-sensing systems, and laying the foundation for novel engineered redox-or gas-sensitive proteins. Such systems could be valuable by both informing aspects of natural biological signaling and in a range of biotech or environmental applications where control of biological processes by redox or heme-binding ligands may be useful.

## Experimental Procedures

### Cloning, Protein Expression and Purification

DNA encoding the FG214 sequence (GenBank: CP007139 Region: 3534516 – 35335172, UNIPROT ID: A0A068NTE8) was ordered from Twist Biosciences and cloned in a pHisGβ1-parallel expression vector(30). FG214 was recombinantly expressed with a His6-Gβ1 tag in BL21(DE3) *E. coli* (Stratagene) in M9 minimal media plus 1 mM 5-aminolevulinic acid at 16°C overnight. Expression was induced with 1 mM isopropyl β-D-1-thiogalactopyranoside. Cells were pelleted and resuspended in Buffer A (50 mM Tris-HCl pH 8.0, 500 mM NaCl). The cells were lysed by sonification, and the lysate was cleared by centrifuging at 10,000 x g for 30 minutes at 4°C. The supernatant was collected and filtered through a 0.2 µm syringe filter. For affinity purification, the protein was loaded onto a 5 mL HisTrap (Cytiva) column preequilibrated with Buffer A. The column was washed with 10 column volumes (CVs) of Buffer A before eluting the protein with buffer B (50mM Tris-HCl pH 8.0, 500 mM NaCl, 500 mM imidazole). The eluent was diluted 1:10 with Buffer C (50 mM Tris-HCl, pH 8.0) and the His_6_-Gβ1 tag was cleaved by adding 1 mg His_6_-TEV protease (70, 71) per 30 mg of fusion protein. After overnight cleavage, the sample was again applied to a preequilibrated 5 mL HisTrap column to remove tag and TEV protease. The flow through was then concentrated using a 30 kDa MWCO Amicon Ultra concentrator to a volume of 3 mL. The concentrated flowthrough was applied to a Superdex 200 10/300 (Cytiva) column preequilibrated with assay buffer (50 mM sodium phosphate pH 7.0 and 50 mM NaCl) unless otherwise noted.

### UV-Visible Absorbance Spectroscopy

FG214 was concentrated to 15 µM in 100 mM sodium phosphate and 50 mM NaCl pH 7.0. Using a 1cm path length quartz cuvette, oxidized spectra were recorded at room temperature using a Varian Cary 60 from 250-600 nm. To reduce the sample, a stock solution of sodium dithionite was made in buffer that previously degassed for 1 hour and added to the sample to a final concentration of 10 mM.

### Mass Spectrometry

To confirm the presence of heme B bound to the protein, we performed a fast SEC-MS experiment using a short SEC column on Dionex Ultimate 3000 LC-system coupled to a Bruker maXis-II ESI-QqTOF mass spectrometer. Briefly, 10 <u>µ</u>L of a 10 µM FG214 stock solution was injected onto a Thermo Scientific NativePac OBE-1 SEC column to enable rapid exchange of non-volatile buffer components into an MS-compatible solvent (50 mM ammonium acetate as the mobile phase). The protein eluted within 2 minutes. We observed the holoprotein with the expected mass increase corresponding to bound heme B, as well as free heme B at m/z 616.18.

### NMR Spectroscopy

^1^H solution-state nuclear magnetic resonance spectra were acquired at 100 µM on a Bruker Avance III HD 800 MHz spectrometer operating at a ^1^H frequency of 800.05 MHz and equipped with a 5mm TCI CryoProbe. All experiments were performed at 298K using the standard Bruker pulse sequence (zgesgp) with NS: 512 and TD: 4096. ^15^N/^1^H TROSY spectra were collected at 250 µM using the pulse sequence ^15^N/^1^H-WADE-TROSY(72) with NS:8 and TD:2048 (F2) x 180 (F1). Oxidized vs. reduced spectra were collected in a valved NMR tube (DWK Life Sciences) to ensure a sustained anaerobic environment. All NMR data were processed and analyzed using NMRFx Analyst(73, 74).

### Hydrogen Deuterium Exchange Mass-Spectrometry

FG214 was concentrated to a final concentration of 40 µM in 100 mM sodium phosphate 50 mM sodium chloride pH 7.0 (with 10 mM sodium dithionite for reduced samples). The protein was diluted 1:14 in 100% D_2_O buffer with same components as the protein buffer, incubated for a defined periods of time at 4°C. The exchange reaction was quenched by adding and equal volume ice-cold quench buffer (3 M guanidine hydrochloride, 3% acetonitrile, and 0.8% formic acid, pH < 2.0), followed by on-line peptic digestion using a Waters Enzymate BEH Pepsin Column, peptide desalting with Hypersil Gold C18 trap column (10 mm) and peptide separation on a Hypersil Gold C18 column (50 mm length × 1 mm diameter, 1.9 μm particle size, Thermo Fisher Scientific). The time between quenching and on-line peptide elution into the mass spectrometer was 5–10 min, at 0–4 °C and pH 2.5, to minimize back-exchange, and peptides were analyzed on a Bruker maXis-II ESI-QqTOF high-resolution mass spectrometer. To minimize in-source back exchange, the instrument ion-source temperature was lowered to 180°C. Initial processing of raw mass spectrometry data files was done with Bruker Compass Data Analysis 6.1. Peptide identification based on accurate mass measurements, MS/MS data, and peptide list curation was performed using Bruker’s Biotools 3.2 and PIGEON(75). Sequence coverage was above 98% for each condition. Downstream data analysis was done using HDExaminer 3.3 (Trajan Scientific). HDExaminer was used for calculation of deuterium uptake profiles and associated standard deviations. Post-analysis quality checks of peptide level deuteration plots, secondary peptide identification and validation were performed using PFNet(75).

### EPR Spectroscopy

Samples were prepared using 200 µL of 250 µM FG214 in assay buffer with 20% glycerol and frozen under liquid nitrogen. Continuous wave X-band EPR spectra were collected on a Bruker EMX spectrometer using Bruker Xenon software (version 1.2) at 10 K using liquid helium and an Oxford Instruments (Oxford, UK) ESR-900 cryogen flow cryostat equipped with an ITC-503 temperature controller. Spectra were simulated using EasySpin 5.2.36 with Matlab R2022(76).

### Size Exclusion Chromatography Multi-Angle Light Scattering

Samples and buffers were filtered with a 0.1 µm pore size filter before use. Samples were injected at onto a Superdex 200 GL 10/300 column pre-equilibrated with 100 mM sodium phosphate and 50 mM NaCl pH 7.0. Following elution light scattering was measured by an inline miniDAWN TREOS and refractive index was measured using an Optilab rEX (Wyatt Technology). The entire process was carried out at room temperature. Data analysis and molecular weight calculations were performed using ASTRA V software (Wyatt Technology).

### Crystallization, Data Processing and X-Ray Structure Determination

Commercially available Morpheus Fusion screen (Molecular Dimensions) was employed to find suitable crystallization conditions. The best diffracting crystals were from condition A4 which contained 0.1 M carboxylic acids mix (0.2 M sodium formate, 0.2 M ammonium acetate, 0.2 M sodium citate tribasic dihydrate, 0.2 M potassium sodium tartrate tetrahydrate and 0.2 M sodium oxamate), 0.12 M alcohols (0.2 M 1,6-hexanediol; 0.2 M 1-butanol; 0.2 M 1,2-propanediol; 0.2 M 2-propanol; 0.2 M 1,4-butanediol and 0.2 M 1,3-propanediol), 0.1 M buffer system 1 at pH 6.5 (0.4 M imidazole and 0.6 M MES monohydrate), and 37.5% precipitant mix 4 (25% v/v MPD, 25% PEG 1000, 25% w/v PEG 3350 w/v). Crystals used for phasing were from condition C7 from the same screen consisting of the following composition, 0.09 M halides (sodium fluoride, sodium bromide and sodium iodide), 0.12 M monosaccharide mix 1 (d-glucose, d-mannose, d-galactose, l-fructose, d-xylose, *N*-acetyl-d-glucosamine) 0.1 M buffer system 1 at pH 6.5 (0.4 M imidazole and 0.6 M MES) and 30% precipitant mix 1 (40% v/v PEG 500 MME and 20% w/v PEG 20000). Crystals were either cryoprotected immediately after opening the well or after a five-minute soak with 10 mM sodium dithionite (Na_2_S_2_O_4_) using LV CryOil (MiTeGen), looped, and flash-cooled in liquid N_2_ prior to data collection. Data were collected at National Synchrotron Light Source II (NSLS-II) light at Brookhaven National Laboratory on beamline 19-ID (NYX). All data sets (C7, A4, and A4 supplemented with Na_2_S_2_O_4_) were measured using 0.920187 Å wavelength and crystallized in space group P2_1_2_1_2_1_ (see Table S2). Data were processed using the autoPROC toolbox(77). Phasing of the A4 data set was achieved through a combination of molecular replacement (MR) with the AlphaFold3 model and single anomalous dispersion (SAD) method using the anomalous signal from the halides present in the crystallization conditions from C7 data set via AutoSol (SAD-MR). Phases for the reduced A4 data set were obtained via MR using the Na_2_S_2_O_4_ free structure. Several cycles of refinement were conducted using Coot(78, 79) and Phenix(80) and further optimization using the PDB_REDO server (Joosten et al. 2014). Final data collection, processing, and refinement parameters are provided in Table S2. The X-ray structure coordinates for truncated FG214 with and without Na_2_S_2_O_4_ are available from the Protein Data Bank under accession codes 10JY and 10JX respectively, with deposited residues 1-3 corresponding to a cloning artifact and 4-145 corresponding to FG214 residues 73-214 in the manuscript.

### Universal Protein Binding Microarrays

Constructs of FG214 wild type and H175I were generated with C-terminal human influenza hemagglutinin (HA) and 6His tags and expressed as described above. Cells were pelleted and resuspended in Buffer A (50 mM Tris-HCl pH 8.0, 500 mM NaCl). The cells were lysed by sonification, and the lysate was cleared by centrifuging at 10,000 x g for 30 minutes at 4°C. The supernatant was collected and filtered through a 0.2 µm syringe filter. For affinity purification, the protein was loaded onto a 5 mL HisTrap (Cytiva) column preequilibrated with Buffer A. The column was washed with 10 column volumes (CVs) of Buffer A before eluting the protein with buffer B (50mM Tris-HCl pH 8.0, 500 mM NaCl, 500 mM imidazole). The eluent was applied to a Superdex 200 10/300 (Cytiva) column preequilibrated with assay buffer (50 mM sodium phosphate pH 7.0 and 50 mM NaCl).

Proteins were assayed simultaneously on separate chambers of one universal PBM (AMADID# 030236, Agilent Technologies, Inc.) as previously described(42) at a final concentration of 750 nM in 1X EN (138 mM potassium glutamate, pH 7.2, 12 mM sodium bicarbonate, 0.8 mM magnesium chloride) or 1X PBS, with or without the inclusion of 500 mM imidazole. PBM scans were analyzed using the Universal PBM Analysis Suite (43) to generate position weight matrices (PWM) and calculate E-scores and Median Signal Intensity for all 8-mers. The representative sequence logo in Figure 5a was generated from the wild-type FG214 PBS+imidazole PWM using the seqLogo package in R. Intensity distributions in Figure 5b show the Median Signal Intensities of 8mers with E-score greater than 0.4, representing the most preferred binding sites, for both alleles in EN.

### Fluorescence Polarization

To create each construct used in the FP assay, FAM labeled ssDNA oligos were synthesized by IDT and annealed to unlabeled reverse compliment ssDNA by applying a decreasing temperature gradient starting at 90°C. Assays were conducted with a constant 1 nM concentration of FAM-labeled dsDNA and increasing concentration of FG214 in the presence of 5 mM MgCl_2_. Imidazole was added to final concentration of 10 µM. Readings were recorded in all black 96-well plates and measured with a Spectramax I3 (Molecular Devices) equipped with a fluorescence polarization cartridge (Molecular Devices) with excitation/emission wavelengths at 485/525 nm. Data reduction was accomplished within the SoftMax software (Molecular Devices).

### Bacterial Two-Hybrid Assays

FG214 constructs were cloned into the multiple cloning sites, attaching them N-terminal of the T18 and T25 fragments of the split adenylyl cyclase using the pUT18 and pKNT25 vectors from the BACTH System Kit(47). Plasmids were transformed into BTH101 cells(47) and grown on plates containing LB-Media, 1.5% agar, 50 µg/mL ampicillin, 50 µg/mL streptomycin, 50 µg/mL kanamycin, 120 µg/mL Bluo-gal, and 0.5 mM IPTG at 30°C for 72 hours. Single colonies were picked and grown overnight in 96 well plates containing 0.5 mL of LB-Media with 50 µg/mL streptomycin, 50 µg/mL kanamycin, 50 µg/mL ampicillin and 0.5 mM IPTG at 30°C. Cultures were diluted 5x in M63 medium (15 mM (NH_4_)_2_SO_4_, 100 mM KH_2_PO_4_, 1.8 µM FeSO_4_, and 1mM MgSO_4_, 0.2% maltose) in 5 mL glass tubes. OD_600_ was recorded using a Spectramax i3 taking 200 µL of the dilution into a 96 well plate. 2.5 mL cultures were permeabilized by adding 30 µL of a 0.1% SDS solution and 30 µL of toluene. After vigorous mixing, the tubes were left in an incubator at 37°C for 40 minutes to allow for toluene evaporation. The enzymatic reaction was started by transferring 500 µL of permeabilized cells to a fresh glass tube containing 500 µL of PM2 buffer (70 mM Na_2_HPO_4_, 30 mM NaH_2_PO_4_, 1 mM MgSO_4_, 0.2 mM MnSO_4_, 100 mM β-mercaptoethanol, and 0.1% *O*-nitrophenol-β-d-galactoside (ONPG) pH 7.0) and allowing reaction to proceed at 28°C for approximately 2 hours. Control wells were made by adding M63 media to the PM2 buffer. The reaction was stopped by addition of 0.5 M Na_2_CO_3_. OD_420_ was recorded and activity was measured by calculating:

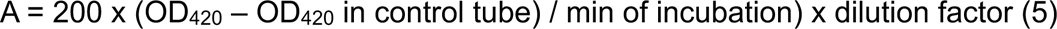

One unit of β-galactosidase activity corresponds to 1 nmol of ONPG hydrolyzed/min at 28°C. The factor 200 is the inverse of the absorption coefficient of the o-nitrophenol, which is 0.005 per nmol/mL at pH 11 (after addition of Na_2_CO_3_). To account for differences in cell density, activity is reported in units/mg dry weight bacteria. OD_600_ was used as 1 mL of culture at OD_600_ = 1 corresponds to 300 µg dry weight bacteria (81, 82). A Welch’s t-test was used to calculate statistical significance of differences in β-galactosidase expression between different samples.

## Data Availability

The X-ray structure coordinates for truncated FG214 with and without Na_2_S_2_O_4_ are available from the Protein Data Bank under accession codes 10JY and 10JX respectively. PBM data are deposited in the NIH Gene Expression Omnibus (GEO) under accession GSE319048. All other data are presented in the manuscript.

## Supporting Information

This article contains supporting information.

## Acknowledgements

We thank members of the Gardner lab as well as Prof. Elizabeth Boon and Jason Withorn (Stony Brook University) for helpful discussions. We thank Prof. Anum Glasgow (Columbia University) and members of her lab assistance running PIGEON and PFNet for HDX-MS peptide list curation and validation, as well as Dr. Paul H. Oyala (Caltech) for EPR access and advice. This work was supported by grants from the NIH (R01 GM106239 and R35 GM156296 to K.H.G.; UC Davis start-up funds for A.H.F.). Use of the NYX beamline (19-ID) at the National Synchrotron Light Source II (NSLS II) is supported by the New York Structural Biology Center. NSLS II is a U.S. Department of Energy (DOE) Office of Science User Facility operated for the DOE Office of Science by Brookhaven National Laboratory under contract DE-SC0012704. This manuscript is the result of funding in whole or in part by the National Institutes of Health (NIH). The content is solely the responsibility of the authors and does not necessarily represent the official views of the National Institutes of Health. It is subject to the NIH Public Access Policy. Through acceptance of this federal funding, NIH has been given a right to make this manuscript publicly available in PubMed Central upon the Official Date of Publication, as defined by NIH.

## References

1. Ulrich, L. E., Koonin, E. V., and Zhulin, I. B. (2005) One-Component Systems Dominate Signal Transduction in Prokaryotes. Trends in Microbiology 13, 52–56.

2. Henry, J. T., and Crosson, S. (2011) Ligand-Binding PAS Domains in a Genomic, Cellular, and Structural Context. Annual Review of Microbiology 65, 261–286.

3. Taylor, B. L., and Zhulin, I. B. (1999) PAS Domains: Internal Sensors of Oxygen, Redox Potential, and Light. Microbiology and Molecular Biology Reviews 63, 479–506.

4. Stuffle, E. C., Johnson, M. S., and Watts, K. J. (2021) PAS Domains in Bacterial Signal Transduction. Current Opinion in Microbiology 61, 8–15.

5. Gilles-Gonzalez, M.-A., and Gonzalez, G. (2004) Signal Transduction by Heme-Containing PAS-Domain Proteins. Journal of Applied Physiology 96, 774–783.

6. Gallio, A. E., Marson, N. A., Heesom, K. J., Lewis, P. A., Alibhai, D., Dugdale, C. A. et al. (2025) An extended network for regulation of heme homeostasis in cells. Proc Natl Acad Sci U S A 122, e2508237122.

7. Farhana, A., Saini, V., Kumar, A., Lancaster, J. R., and Steyn, A. J. C. (2012) Environmental Heme-Based Sensor Proteins: Implications for Understanding Bacterial Pathogenesis. Antioxidants & Redox Signaling 17, 1232–1245.

8. Taabazuing, C. Y., Hangasky, J. A., and Knapp, M. J. (2014) Oxygen Sensing Strategies in Mammals and Bacteria. Journal of Inorganic Biochemistry 133, 63–72.

9. Chim, N., Johnson, P. M., and Goulding, C. W. (2014) Insights into Redox Sensing Metalloproteins in Mycobacterium tuberculosis. Journal of Inorganic Biochemistry 133, 118–126.

10. Nash, A. I., McNulty, R., Shillito, M. E., Swartz, T. E., Bogomolni, R. A., Luecke, H. et al. (2011) Structural Basis of Photosensitivity in a Bacterial Light-Oxygen-Voltage/Helix-Turn-Helix (LOV-HTH) DNA-Binding Protein. Proceedings of the National Academy of Sciences 108, 9449–9454.

11. Rivera-Cancel, G., Motta-Mena, L. B., and Gardner, K. H. (2012) Identification of Natural and Artificial DNA Substrates for Light-Activated LOV–HTH Transcription Factor EL222. Biochemistry 51, 10024–10034.

12. Zoltowski, B. D., Motta-Mena, L. B., and Gardner, K. H. (2013) Blue Light-Induced Dimerization of a Bacterial LOV–HTH DNA-Binding Protein. Biochemistry 52, 6653–6661.

13. Chaudhari, A. S., Favier, A., Tehrani, Z. A., Koval, T., Andersson, I., Schneider, B. et al. (2025) Light-dependent flavin redox and adduct states control the conformation and DNA-binding activity of the transcription factor EL222. Nucleic Acids Res 53, gkaf215.

14. Motta-Mena, L. B., Reade, A., Mallory, M. J., Glantz, S., Weiner, O. D., Lynch, K. W. et al. (2014) An Optogenetic Gene Expression System with Rapid Activation and Deactivation Kinetics. Nature Chemical Biology 10, 196–202.

15. Reade, A., Motta-Mena, L. B., Gardner, K. H., Stainier, D. Y., Weiner, O. D., and Woo, S. (2016) TAEL: A Zebrafish-Optimized Optogenetic Gene Expression System with Fine Spatial and Temporal Control. Development 144, 345–355.

16. Rullan, M., Benzinger, D., Schmidt, G. W., Milias-Argeitis, A., and Khammash, M. (2018) An Optogenetic Platform for Real-Time, Single-Cell Interrogation of Stochastic Transcriptional Regulation. Molecular Cell 70, 745–756.e746.

17. Zhao, E. M., Lalwani, M. A., Lovelett, R. J., García-Echauri, S. A., Hoffman, S. M., Gonzalez, C. L. et al. (2020) Design and Characterization of Rapid Optogenetic Circuits for Dynamic Control in Yeast Metabolic Engineering. ACS Synthetic Biology 9, 3254–3266.

18. Zhao, E. M., Zhang, Y., Mehl, J., Park, H., Lalwani, M. A., Toettcher, J. E. et al. (2018) Optogenetic Regulation of Engineered Cellular Metabolism for Microbial Chemical Production. Nature 555, 683–687.

19. Cleere, M. M., and Gardner, K. H. (2024) Optogenetic Control of Phosphate-Responsive Genes Using Single-Component Fusion Proteins in Saccharomyces cerevisiae. ACS Synthetic Biology 13, 4085–4098.

20. Hoffman, S. M., Espinel-Rios, S., Kwartler, S. K., Lalwani, M. A., and Avalos, J. L. (2025) Balancing Doses of EL222 and Light Improves Optogenetic Induction of Protein Production in Komagataella phaffii. Biotechnol Bioeng 122, 2478–2498.

21. Im, W.-T., Hu, Z.-Y., Kim, K.-H., Rhee, S.-K., Meng, H., Lee, S.-T. et al. (2012) Description of Fimbriimonas ginsengisoli gen. nov., sp. nov. within the Fimbriimonadia Class nov., of the Phylum Armatimonadetes. Antonie van Leeuwenhoek 102, 307–317.

22. Siclari, J. J., Favaro, D. C., Huang, R. H., and Gardner, K. H. (2025) A pipeline for screening small molecule-enhanced protein stability in a bacterial orphan receptor. Protein Sci 34, e70330.

23. Letunic, I., Khedkar, S., and Bork, P. (2021) SMART: Recent Updates, New Developments and Status in 2020. Nucleic Acids Research 49, D458–D460.

24. Wißbrock, A., Paul George, A. A., Hans, Kühl, T., and Imhof, D. (2019) The Molecular Basis of Transient Heme-Protein Interactions: Analysis, Concept and Implementation. Bioscience Reports 39, BSR20181940.

25. San Francisco, B., Bretsnyder, E. C., and Kranz, R. G. (2013) Human Mitochondrial Holocytochrome C Synthase’s Heme Binding, Maturation Determinants, and Complex Formation with Cytochrome C. Proceedings of the National Academy of Sciences 110, E788–E797.

26. Ghosh, K., Thompson, A. M., Goldbeck, R. A., Shi, X., Whitman, S., Oh, E. et al. (2005) Spectroscopic and Biochemical Characterization of Heme Binding to Yeast Dap1p and Mouse PGRMC1p. Biochemistry 44, 16729–16736.

27. Ishitsuka, Y., Araki, Y., Tanaka, A., Igarashi, J., Ito, O., and Shimizu, T. (2008) Arg97 at the Heme-Distal Side of the Isolated Heme-Bound PAS Domain of a Heme-Based Oxygen Sensor from Escherichia coli (Ec DOS) Plays Critical Roles in Autoxidation and Binding to Gases, Particularly O2. Biochemistry 47, 8874–8884.

28. Bondarenko, V., Dewilde, S., Moens, L., and La Mar, G. N. (2006) Solution 1H NMR characterization of the axial bonding of the two His in oxidized human cytoglobin. J Am Chem Soc 128, 12988–12999.

29. Gochin, M., and Roder, H. (1995) Protein Structure Refinement Based on Paramagnetic NMR Shifts: Applications to Wild-Type and Mutant Forms of Cytochrome c. Protein Science 4, 296–305.

30. Harper, S. M., Neil, L. C., and Gardner, K. H. (2003) Structural Basis of a Phototropin Light Switch. Science 301, 1541–1544.

31. Yao, X., Rosen, M. K., and Gardner, K. H. (2008) Estimation of the Available Free Energy in a LOV2-Jα Photoswitch. Nature Chemical Biology 4, 491–497.

32. Imelio, J. A., Trajtenberg, F., and Buschiazzo, A. (2021) Allostery and Protein Plasticity: the Keystones for Bacterial Signaling and Regulation. Biophysical Reviews 13, 943–953.

33. Kiel, C., Matallanas, D., and Kolch, W. (2021) The Ins and Outs of RAS Effector Complexes. Biomolecules 11, 236.

34. Zhang, N., Dong, Y., Zhou, H., and Cui, H. (2022) Effect of PAS-LuxR Family Regulators on the Secondary Metabolism of Streptomyces. Antibiotics 11, 1783.

35. Li, T., Bonkovsky, H. L., and Guo, J.-T. (2011) Structural Analysis of Heme Proteins: Implications for Design and Prediction. BMC Structural Biology 11, 13.

36. Zoppellaro, G., Bren, K. L., Ensign, A. A., Harbitz, E., Kaur, R., Hersleth, H. P. et al. (2009) Review: Studies of Ferric Heme Proteins with Highly Anisotropic/Highly Axial Low Spin (S = 1/2) Electron Paramagnetic Resonance Signals with Bis-Histidine and Histidine-Methionine Axial Iron Coordination. Biopolymers 91, 1064–1082.

37. Tanaka, A., and Shimizu, T. (2008) Ligand Binding to the Fe(III)-Protoporphyrin IX Complex of Phosphodiesterase from Escherichia coli (EcDOS) Markedly Enhances Catalysis of Cyclic di-GMP: Roles of Met95, Arg97, and Phe113 of the Putative Heme Distal Side in Catalytic Regulat. Biochemistry 47, 13438–13446.

38. Fojtikova, V., Stranava, M., Vos, M. H., Liebl, U., Hranicek, J., Kitanishi, K. et al. (2015) Kinetic Analysis of a Globin-Coupled Histidine Kinase, Af GcHK: Effects of the Heme Iron Complex, Response Regulator, and Metal Cations on Autophosphorylation Activity. Biochemistry 54, 5017–5029.

39. Surette, M. G., and Stock, J. B. (1996) Role of α-Helical Coiled-Coil Interactions in Receptor Dimerization, Signaling, and Adaptation during Bacterial Chemotaxis. Journal of Biological Chemistry 271, 17966–17973.

40. Macheboeuf, P., Buffalo, C., Fu, C.-Y., Zinkernagel, A. S., Cole, J. N., Johnson, J. E. et al. (2011) Streptococcal M1 Protein Constructs a Pathological Host Fibrinogen Network. Nature 472, 64–68.

41. Hurme, R., Berndt, K. D., Namork, E., and Rhen, M. (1996) DNA Binding Exerted by a Bacterial Gene Regulator with an Extensive Coiled-coil Domain. Journal of Biological Chemistry 271, 12626–12631.

42. Berger, M. F., Philippakis, A. A., Qureshi, A. M., He, F. S., Estep, P. W., and Bulyk, M. L. (2006) Compact, Universal DNA Microarrays to Comprehensively Determine Transcription-Factor Binding Site Specificities. Nature Biotechnology 24, 1429–1435.

43. Berger, M. F., and Bulyk, M. L. (2009) Universal Protein-Binding Microarrays for the Comprehensive Characterization of the DNA-Binding Specificities of Transcription Factors. Nature Protocols 4, 393–411.

44. Bidwai, A. K., Ahrendt, A. J., Sullivan, J. S., Vitello, L. B., and Erman, J. E. (2015) pH Dependence of Cyanide and Imidazole Binding to the Heme Domains of Sinorhizobium meliloti and Bradyrhizobium japonicum FixL. Journal of Inorganic Biochemistry 153, 88–102.

45. Huang, X., and Groves, J. T. (2018) Oxygen Activation and Radical Transformations in Heme Proteins and Metalloporphyrins. Chemical Reviews 118, 2491–2553.

46. Gong, W., Hao, B., and Chan, M. K. (2000) New Mechanistic Insights from Structural Studies of the Oxygen-Sensing Domain of Bradyrhizobium japonicum FixL. Biochemistry 39, 3955–3962.

47. Olson, M. G., Goldammer, M., Gauliard, E., Ladant, D., and Ouellette, S. P. (2018) A Bacterial Adenylate Cyclase-Based Two-Hybrid System Compatible with Gateway® Cloning Springer New York, 75–96

48. Karimova, G., Dautin, N., and Ladant, D. (2005) Interaction Network among Escherichia coli Membrane Proteins Involved in Cell Division as Revealed by Bacterial Two-Hybrid Analysis. Journal of Bacteriology 187, 2233–2243.

49. Möglich, A., Ayers, R. A., and Moffat, K. (2009) Structure and Signaling Mechanism of Per-ARNT-Sim Domains. Structure 17, 1282–1294.

50. Glantz, S. T., Carpenter, E. J., Melkonian, M., Gardner, K. H., Boyden, E. S., Wong, G. K.-S. et al. (2016) Functional and Topological Diversity of LOV Domain Photoreceptors. Proceedings of the National Academy of Sciences 113, E1442–E1451.

51. Chawla, M., Parikh, P., Saxena, A., Munshi, M., Mehta, M., Mai, D. et al. (2012) Mycobacterium tuberculosis WhiB4 Regulates Oxidative Stress Response to Modulate Survival and Dissemination in vivo. Molecular Microbiology 85, 1148–1165.

52. Mondragon, P., Hwang, S., Kasirajan, L., Oyetoro, R., Nasthas, A., Winters, E. et al. (2022) TrmB Family Transcription Factor as a Thiol-Based Regulator of Oxidative Stress Response. mBio 13, e0063322.

53. Li, Y., and He, Z.-G. (2012) The Mycobacterial LysR-Type Regulator OxyS Responds to Oxidative Stress and Negatively Regulates Expression of the Catalase-Peroxidase Gene. PLoS ONE 7, e30186.

54. Banerjee, S., Chanakira, M. N., Hall, J., Kerkan, A., Dasgupta, S., and Martin, D. W. (2022) A Review on Bacterial Redox Dependent Iron Transporters and their Evolutionary Relationship. Journal of Inorganic Biochemistry 229, 111721.

55. Ciemniecki, J. A., and Newman, D. K. (2020) The Potential for Redox-Active Metabolites To Enhance or Unlock Anaerobic Survival Metabolisms in Aerobes. J Bacteriol 202, 10.1128/JB.00797-00719.

56. Monson, E. K., Weinstein, M., Ditta, G. S., and Helinski, D. R. (1992) The FixL Protein of Rhizobium meliloti can be Separated into a Heme-Binding Oxygen-Sensing Domain and a Functional C-Terminal Kinase Domain. Proceedings of the National Academy of Sciences 89, 4280–4284.

57. Orillard, E., Anaya, S., Johnson, M. S., and Watts, K. J. (2021) Oxygen-Induced Conformational Changes in the PAS-Heme Domain of the Pseudomonas aeruginosa Aer2 Receptor. Biochemistry 60, 2610–2622.

58. Delgado-Nixon, V. M., Gonzalez, G., and Gilles-Gonzalez, M.-A. (2000) Dos, a Heme-Binding PAS Protein from Escherichia coli, Is a Direct Oxygen Sensor. Biochemistry 39, 2685–2691.

59. Garcia, D., Orillard, E., Johnson, M. S., and Watts, K. J. (2017) Gas Sensing and Signaling in the PAS-Heme Domain of the Pseudomonas aeruginosa Aer2 Receptor. Journal of Bacteriology 199, JB.00003–00017.

60. Dent, M. R., DeMartino, A. W., Xu, Q., Chen, X., Gandhi, A., Hwang, H. S. et al. (2025) Engineering a highly selective, hemoprotein-based scavenger as a carbon monoxide poisoning antidote with no hypertensive effect. Proc Natl Acad Sci U S A 122, e2501389122.

61. Kerby, R. L., Youn, H., and Roberts, G. P. (2008) RcoM: A New Single-Component Transcriptional Regulator of CO Metabolism in Bacteria. Journal of Bacteriology 190, 3336–3343.

62. Mukherjee, P., Agarwal, S., Mallick, S. B., and Dasgupta, J. (2024) PAS domain of flagellar histidine kinase FlrB has a unique architecture and binds heme as a sensory ligand in an unconventional fashion. Structure 32, 200–216.e205.

63. Noya, F., Arias, A., and Fabiano, E. (1997) Heme Compounds as Iron Sources for Nonpathogenic Rhizobium Bacteria. Journal of Bacteriology 179, 3076–3078.

64. Zygiel, E. M., Obisesan, A. O., Nelson, C. E., Oglesby, A. G., and Nolan, E. M. (2021) Heme protects Pseudomonas aeruginosa and Staphylococcus aureus from calprotectin-induced iron starvation. J Biol Chem 296, 100160.

65. Gong, W., Hao, B., Mansy, S. S., Gonzalez, G., Gilles-Gonzalez, M. A., and Chan, M. K. (1998) Structure of a Biological Oxygen Sensor: A New Mechanism for Heme-Driven Signal Transduction. Proceedings of the National Academy of Sciences 95, 15177–15182.

66. Reynolds, M. F. (2024) New Insights into the Signal Transduction Mechanism of O2-Sensing FixL and Other Biological Heme-Based Sensor Proteins. Journal of Inorganic Biochemistry 259, 112642.

67. Mokdad, A., Ang, E., Desciak, M., Ott, C., Vilbert, A., Beddow, O. et al. (2024) Photoacoustic Calorimetry Studies of O2-Sensing FixL and (R200, I209) Variants from Sinorhizobium meliloti Reveal Conformational Changes Coupled to Ligand Photodissociation from the Heme-PAS Domain. Biochemistry 63, 116–127.

68. Wright, G. S. A., Saeki, A., Hikima, T., Nishizono, Y., Hisano, T., Kamaya, M. et al. (2018) Architecture of the Complete Oxygen-Sensing FixL-FixJ Two-Component Signal Transduction System. Science Signaling 11, eaaq0825.

69. Guimarães, W. G., Lobão, J. B. D. S., Carepo, M. S. P., Gondim, A. C. S., Lopes, L. G. D. F., Paulo, T. D. F. et al. (2026) Circular Dichroism: A Possible Route to Shed Light on Signal Transduction Processes of Heme-Based Gas-Sensor Proteins. Journal of Inorganic Biochemistry 280, 113311.

70. Kapust, R. B., Tözsér, J., Fox, J. D., Anderson, D. E., Cherry, S., Copeland, T. D. et al. (2001) Tobacco etch virus protease: mechanism of autolysis and rational design of stable mutants with wild-type catalytic proficiency. Protein Engineering, Design and Selection 14, 993–1000.

71. Phan, J., Zdanov, A., Evdokimov, A. G., Tropea, J. E., Peters, H. K., Kapust, R. B. et al. (2002) Structural Basis for the Substrate Specificity of Tobacco Etch Virus Protease. Journal of Biological Chemistry 277, 50564–50572.

72. Manu, V. S., Olivieri, C., Pavuluri, K., and Veglia, G. (2022) Design and applications of water irradiation devoid RF pulses for ultra-high field biomolecular NMR spectroscopy. Physical Chemistry Chemical Physics 24, 18477–18481.

73. Norris, M., Fetler, B., Marchant, J., and Johnson, B. A. (2016) NMRFx Processor: a cross-platform NMR data processing program. Journal of Biomolecular NMR 65, 205–216.

74. Koag, E., Hulse, S. G., Helms, G. L., Call, K. M., Summers, M. F., Marchant, J. et al. (2025) NMRFx: Integrated Software for NMR Data Processing, Visualization, Analysis and Structure Calculation Cold Spring Harbor Laboratory,

75. Lu, C., Wells, M. L., Reckers, A., McBride, S. K., and Glasgow, A. (2026) Site-resolved energetic information from HX-MS experiments. Nat Chem Biol 22, 307–317.

76. Stoll, S., and Schweiger, A. (2006) EasySpin, a comprehensive software package for spectral simulation and analysis in EPR. Journal of Magnetic Resonance 178, 42–55.

77. Vonrhein, C., Flensburg, C., Keller, P., Sharff, A., Smart, O., Paciorek, W., et al. (2011) Data processing and analysis with the *autoPROC*toolbox. Acta Crystallographica Section D Biological Crystallography 67, 293–302.

78. Agirre, J., Atanasova, M., Bagdonas, H., Ballard, C. B., Baslé, A., Beilsten-Edmands, J., et al. (2023) The *CCP*4 suite: integrative software for macromolecular crystallography. Acta Crystallographica Section D Structural Biology 79, 449–461.

79. Emsley, P., Lohkamp, B., Scott, W. G., and Cowtan, K. (2010) Features and development of *Coot*. Acta Crystallographica Section D Biological Crystallography 66, 486–501.

80. Liebschner, D., Afonine, P. V., Baker, M. L., Bunkóczi, G., Chen, V. B., Croll, T. I., et al. (2019) Macromolecular structure determination using X-rays, neutrons and electrons: recent developments in *Phenix*. Acta Crystallographica Section D Structural Biology 75, 861–877.

81. Karimova, G., Dautin, N., and Ladant, D. (2005) Interaction Network among *Escherichia coli*Membrane Proteins Involved in Cell Division as Revealed by Bacterial Two-Hybrid Analysis. Journal of Bacteriology 187, 2233–2243.

82. Hosseini, A., and Mas, J. (2021) The β-galactosidase assay in perspective: Critical thoughts for biosensor development. Analytical Biochemistry 635, 114446.

