## Supporting Information for "Structural Basis of a Novel Heme Binding Bacterial One-Component Switch"

James J. Siclari[1,2], Malvin Forson[1,3], Cullen Roeder[1,3], Eta A. Isiorho[1], Denize C. Favaro[1], Rinat R. Abzalimov[1], Stephen S. Gisselbrecht[4], Alec H. Follmer[5], Martha L. Bulyk[4,6], Kevin H. Gardner[1,7,8,*]

[1]: Structural Biology Initiative, CUNY Advanced Science Research Center, New York, NY 10031
[2]: Ph.D. Program in Biology, The Graduate Center – City University of New York, New York, NY 10016
[3]: Ph.D. Program in Biochemistry, The Graduate Center – City University of New York, New York, NY 10016
[4]: Division of Genetics, Department of Medicine, Brigham and Women's Hospital and Harvard Medical School, Boston, MA 02115
[5]: Department of Chemistry, University of California Davis, Davis, CA 95616, USA
[6]: Department of Pathology, Brigham and Women's Hospital and Harvard Medical School, Boston, MA 02115
[7]: Ph.D. Programs in Biochemistry, Biology, and Chemistry, The Graduate Center – City University of New York, New York, NY 10016
[8]: Department of Chemistry and Biochemistry, City College of New York, New York, NY 10031

Contents:

- Supporting Figures S1-S6
- Supporting Tables S1 & S2

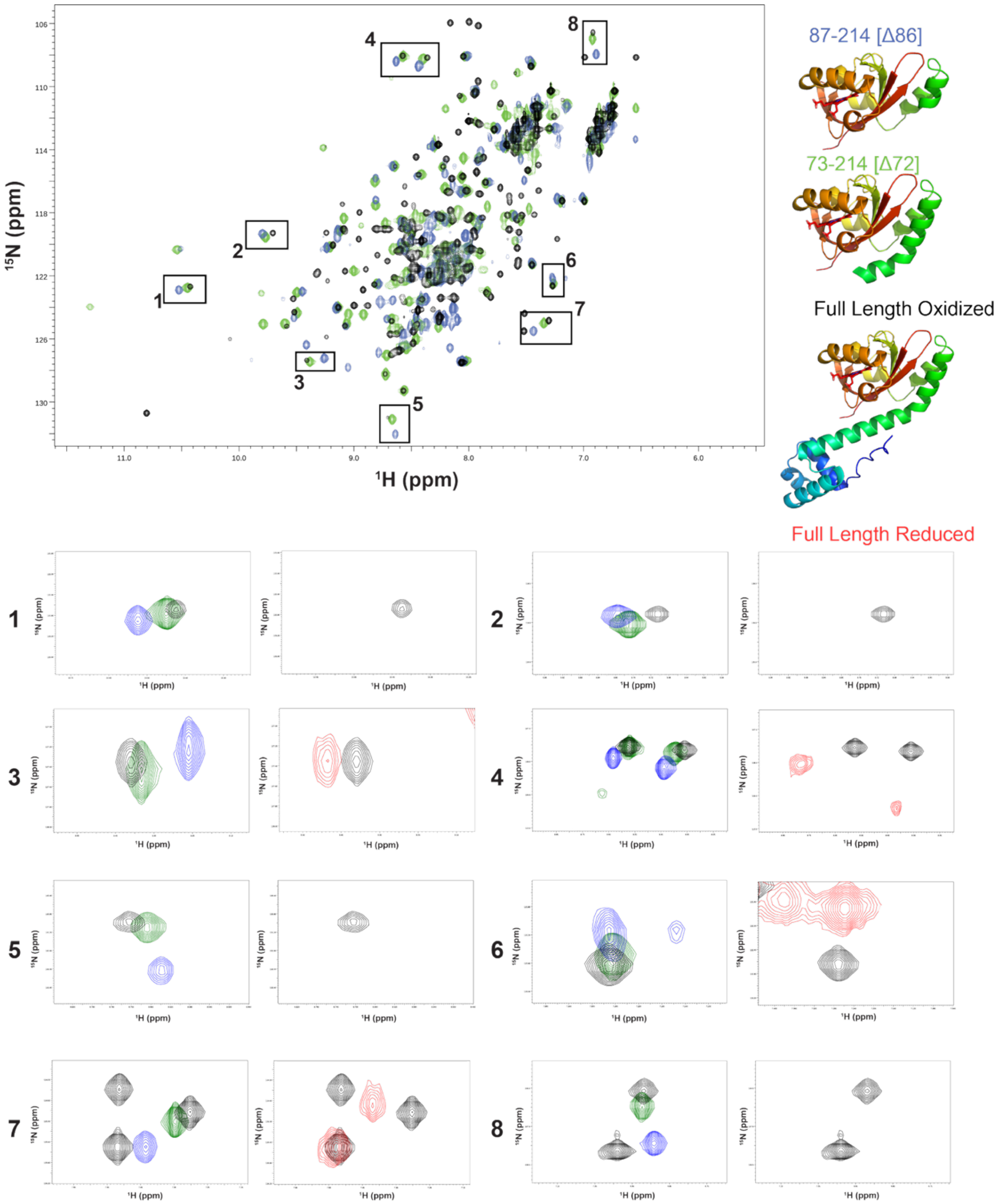

**<u>Figure S1:</u>** $^{15}$N/$^{1}$H HSQC spectra of oxidized FG214 constructs: Full Length (Black), Δ72 (green), and Δ86 (blue). Boxes indicate peaks that display progressive perturbation in all constructs, and insets show expanded regions compared to full length oxidized (black) and full length reduced (red).

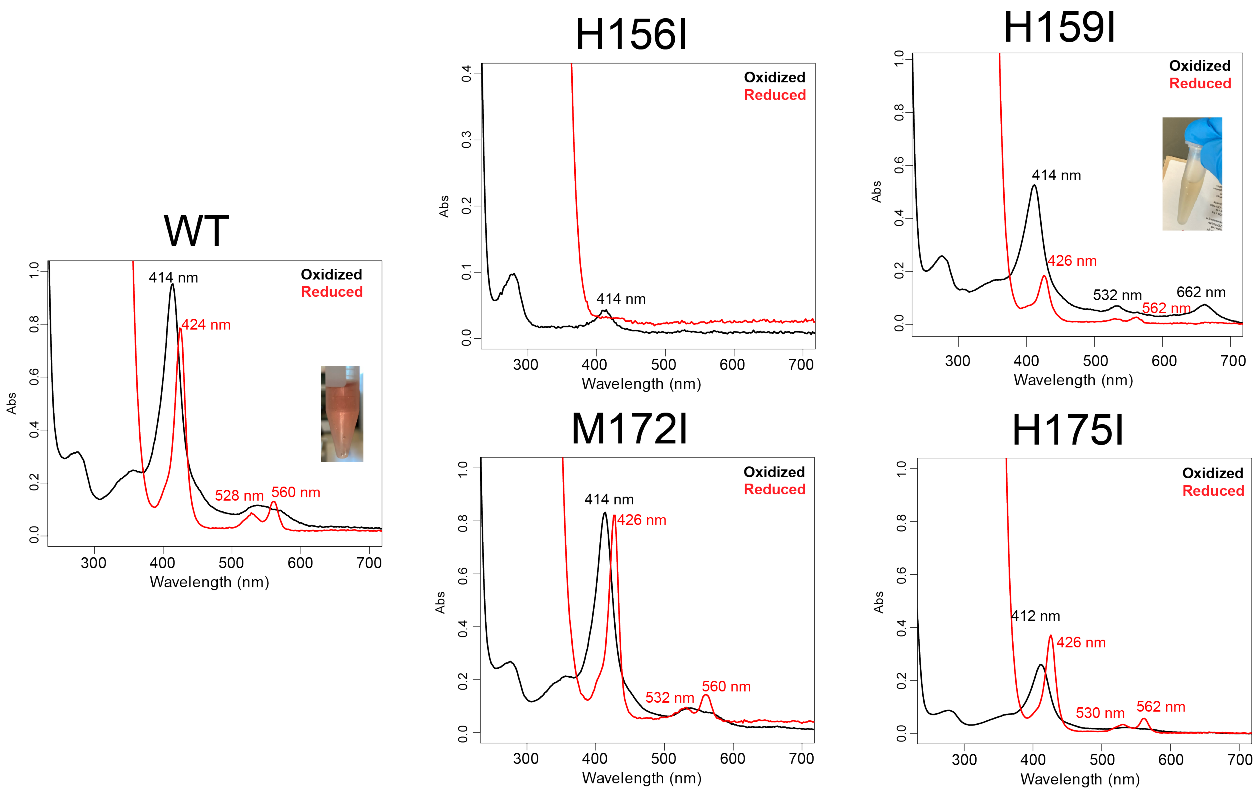

**Figure S2:** UV-Visible absorbance spectra of FG214 wild-type (WT) and PAS domain point mutants under oxidized (black) and reduced (red) conditions. WT data are reproduced here from Figure 1d for ease of comparison.

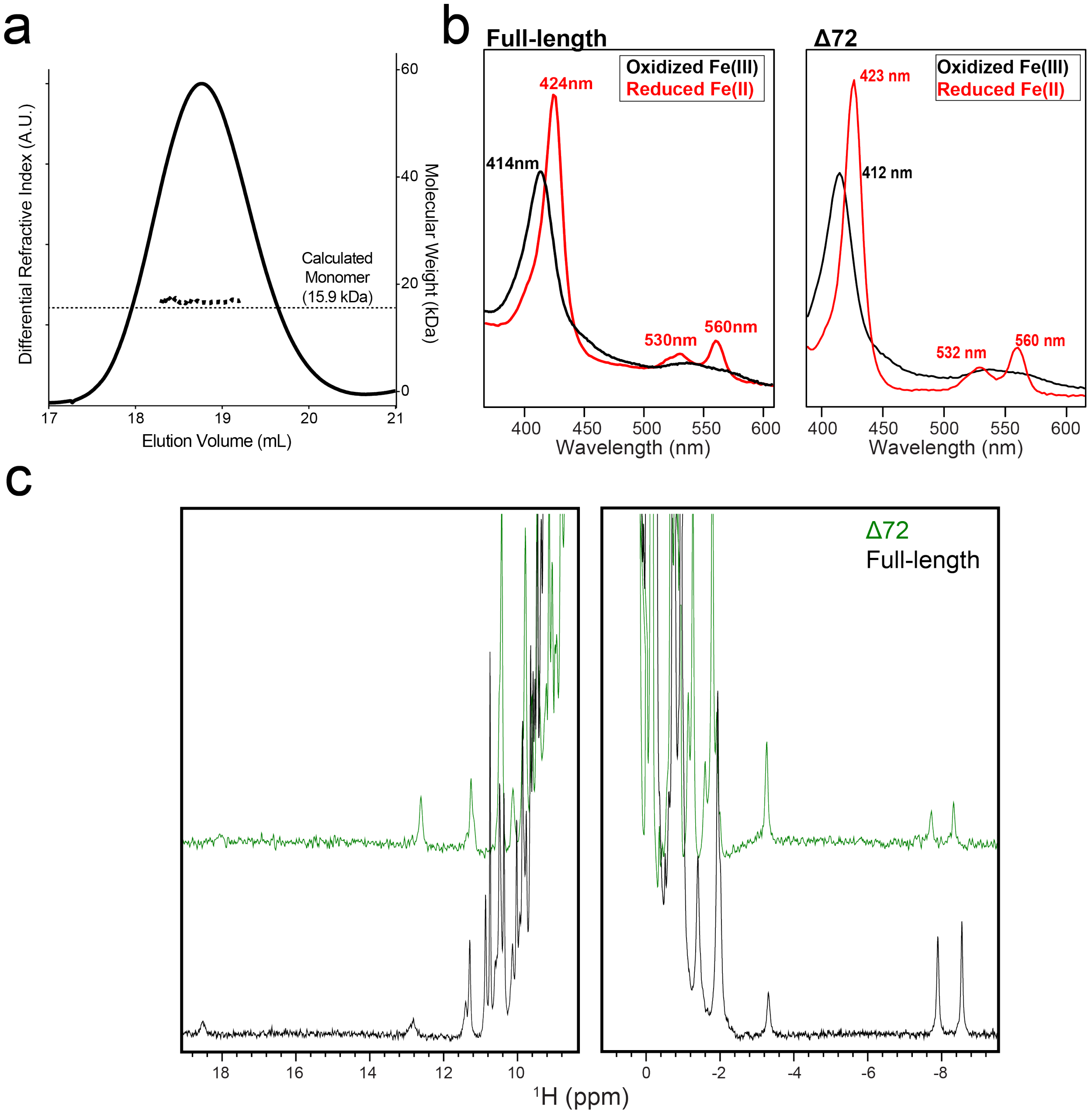

**<u>Figure S3</u>**: Solution characterization of FG214 (Δ72). **a)** SEC-MALS analysis of FG214 (Δ72) compared to calculated monomeric molecular weight. **b)** UV-Visible absorbance spectra of oxidized (black) and reduced (red) FG214 (Δ72) compared to full-length data as shown in Figure 1. **c)** $^1$H NMR spectra of oxidized FG214 (Δ72) (green) compared to full-length (black, reproduced from Figure 1f).

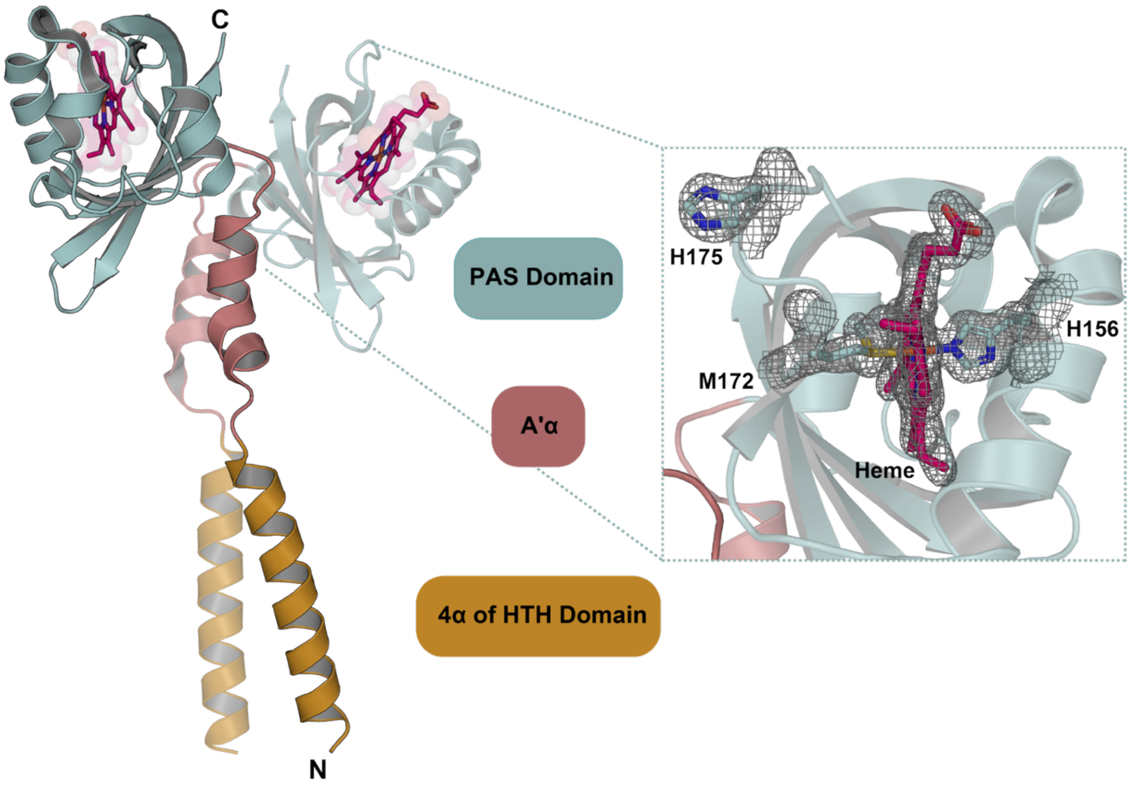

**<u>Figure S4:</u>** 1.65 Å crystal structure of FG214(Δ72) from the same crystal conditions yielding the structure from Fig. 4 but cryoprotected following addition of soluble sodium dithionite. PDB: 10JY.

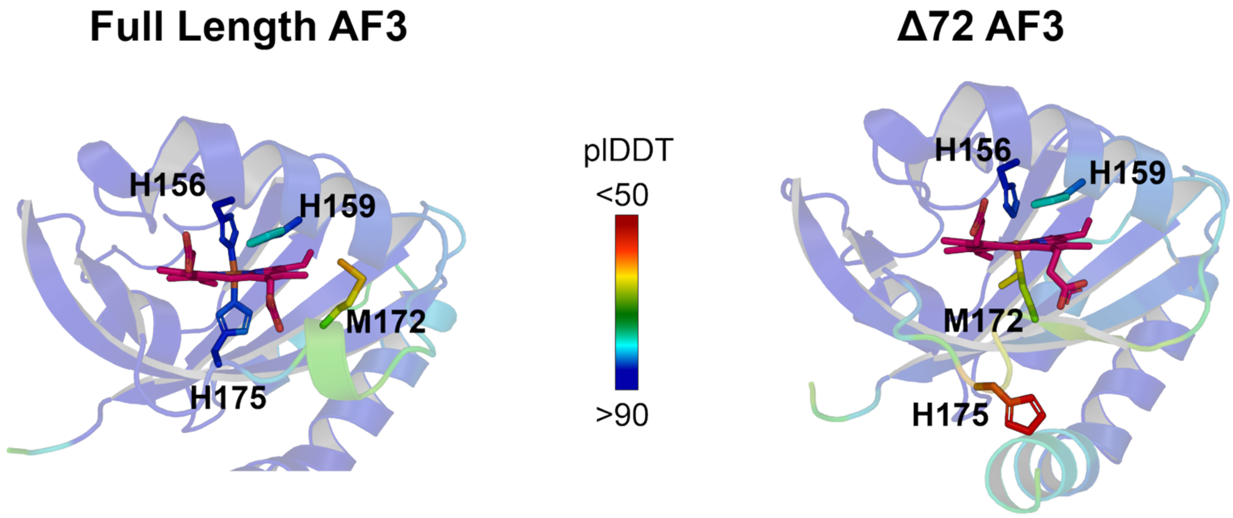

**<u>Figure S5:</u>** AlphaFold3 model of heme binding pocket predicted in full length model (left) compared to Δ72 model (right).

Direct: atat-GGGGCGGGG-(n)-GGGGCGGGG-aatt

Inverted: atat-GGGGCGGGG-(n)-CCCCGCCCC-aatt

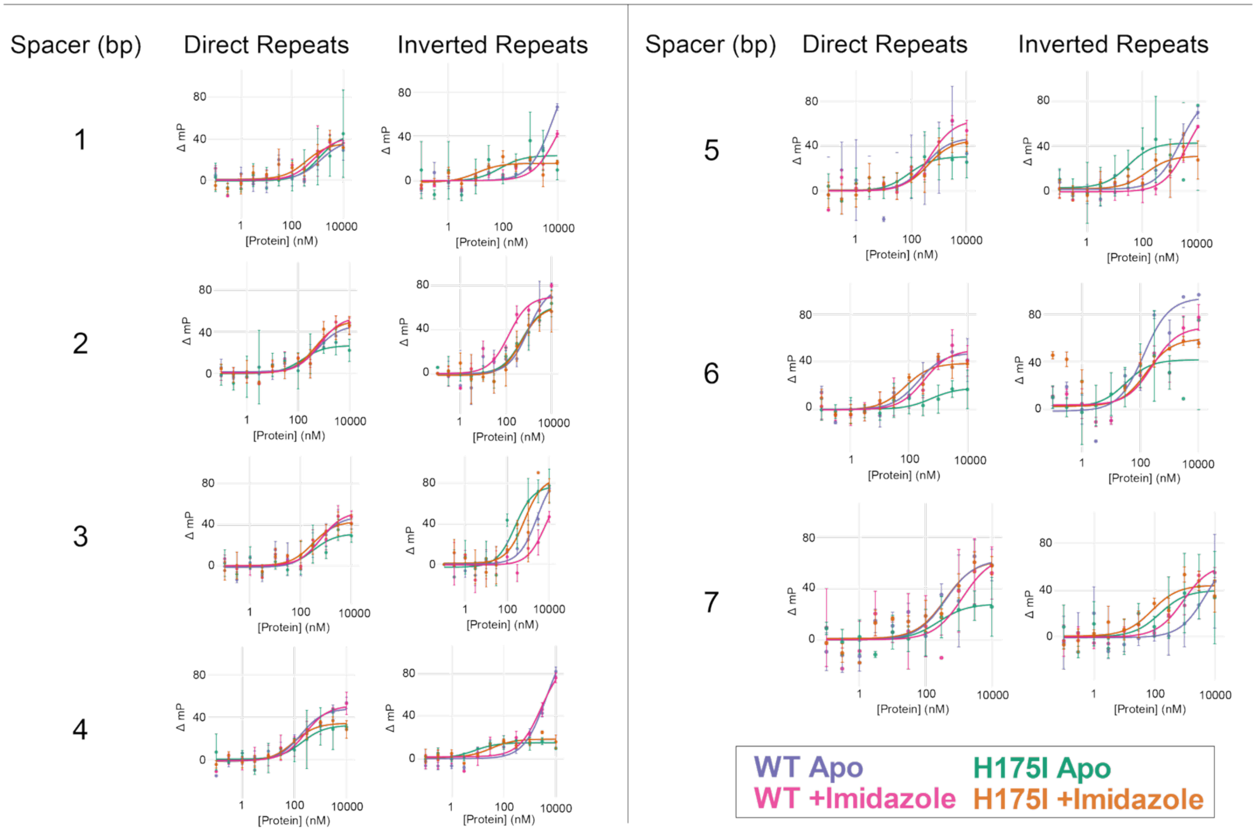

**Figure S6:** Fluorescence polarization analysis collected in triplicate for H175I apo (green), H175I + imidazole (orange), WT apo (purple), and WT + imidazole (pink).

**<u>Table S1:</u> UV-visible absorbance spectra peaks for FG214 constructs in oxidized and reduced conditions**

| Mutant | State | Soret Band (nm) | Q Bands (nm) | Additional Peak (nm) |
|---|---|---|---|---|
| WT | Oxidized | 414 | Broad 550 | Not observed |
| | Reduced | 424 | 528 / 560 | Not observed |
| H156I | Oxidized | 414 | Not observed | Not observed |
| | Reduced | Not observed | Not observed | Not observed |
| H159I | Oxidized | 414 | 532 | 662 |
| | Reduced | 426 | 532 / 562 | Not observed |
| M172I | Oxidized | 414 | Broad 550 | Not observed |
| | Reduced | 427 | 532 / 560 | Not observed |
| H175I | Oxidized | 412 | Broad 550 | Not observed |
| | Reduced | 426 | 530 / 562 | Not observed |

**Table S2: FG214 Crystallographic Data Collection and Refinement Statistics**

| | FG214(73-214) | FG214(73-214)+DT Soak |
|---|---|---|
| Data Collection | | |
| Space Group | $P2_1\ 2_1\ 2_1$ | $P2_1\ 2_1\ 2_1$ |
| Unit cell parameters (Å) | | |
| | a = 63.35 | a = 63.03 |
| | b = 63.93 | b = 65.21 |
| | c = 105.97 | c = 105.87 |
| Wavelength (Å) | 0.92019 | 0.97860 |
| Resolution Range (Å) | 34.451 – 1.467 (1.493 – 1.467) | 55.523 – 1.648 (1.677 – 1.648) |
| Total reflections | 583436 | 393593 |
| Unique reflections | 75338 (3680) | 53371 (2645) |
| Multiplicity | 7.7 (7.0) | 7.4 (7.7) |
| Completeness (%) | 99.9 (99.9) | 100.0 (99.9) |
| Mean $I/\sigma(I)$ | 9.8 (1.5) | 11.0 (0.6) |
| $R_{meas}$ | 0.115 (2.397) | 0.093 (3.327) |
| $CC_{1/2}$ | 0.992 (0.335) | 0.999 (0.382) |
| Refinement | | |
| Reflection used in refinement | 75301 (2823) | 53262 (2716) |
| Reflections used in R-free | 3708 (132) | 2634 (141) |
| Number of non-hydrogen atom | 2656 | 2563 |
| Macromolecules | 2333 | 2286 |
| Ligands | 139 | 105 |
| Solvent | 183 | 172 |
| $R_{work}$ | 0.1641 (0.3044) | 0.1899 (0.3734) |
| $R_{free}$ | 0.1901 (0.2984) | 0.2150 (0.3636) |
| RMS(bonds) | 0.006 | 0.008 |
| RMS(angles) | 0.87 | 0.93 |
| Protein residues | 282 | 286 |
| Average B-factor | 29.1 | 38.67 |
| Macromolecules | 27.99 | 38.58 |
| Ligands | 37.05 | 34.5 |
| Solvent | 36.76 | 42.5 |
| Ramachandran plots | | |
| Favored (%) | 98.55 | 100.0 |
| Allowed (%) | 1.09 | 0.00 |
| Outliers (%) | 0.36 | 0.00 |
| PDB accession code | 10JX | 10JY |

Statistics for the highest-resolution shell are shown in parentheses. Note that residues 4-145 in the PDB entries correspond to FG214 residues 73-214 in the manuscript; PDB residues 1-3 are derived from a cloning artifact.

### Table S3: DNA Sequences used in Figure 5C

| Name | Sequence (5'-3') |
|---|---|
| **IR2** | ATATGGGGCGGGGATCCCCGCCCCAATT |
| **DR2** | ATATGGGGCGGGGATGGGGCGGGGAATT |
